# GTP hydrolysis triggers membrane remodeling by AMPH-1

**DOI:** 10.1101/2024.08.16.608333

**Authors:** Wei Gai, Yuhang Wang, Junjie Zhang, Chavela M. Carr, Hays S. Rye

## Abstract

Membrane-enclosed transport carriers return biological molecules from the recycling endosome to the plasma membrane using a mechanism that is not well understood. In *C. elegans*, the formation of carriers from the recycling endosome requires the amphiphysin protein, AMPH-1. Recently, we found that purified AMPH-1 is sufficient for tubulation and vesiculation of liposomes in a mechanism that is regulated by guanine nucleotides. Here we propose a model linking GTP binding and hydrolysis to the membrane binding and tubulation required for transport carrier formation. We find that GTP binding stabilizes interactions between AMPH-1 and the membrane through amphipathic, N-terminal alpha helices, which are found at the tips of the arc-shaped, homodimeric structure. By contrast, in the post-hydrolysis, GDP-bound state, these helices are repositioned to interact with the N-terminal helices of other homodimers, to form an oligomeric AMPH-1 lattice that tubulates the membrane, in preparation for carrier formation by membrane fission.

## Introduction

Endocytic recycling is the process that returns lipids and membrane proteins to the cell surface following internalization and sorting ^1–3^. Recycling defects have been implicated in a wide range of diseases, such as diabetes ^4,5^, cancer ^6–9^, and mental retardation ^10^. Understanding how endocytic recycling is regulated is expected to facilitate the development of therapeutics designed to mitigate the effects of these recycling defects.

While the formation and release of tubular-vesicular carriers from the plasma membrane ^3,11^ and early endosomes ^12^ have been well characterized, the precise mechanism of carrier formation from recycling endosomes remains unresolved. During endocytosis, proteins of the Epsin N-Terminal Homology (ENTH) domain, Eps15 Homology Domain (EHD), and various Bin/Amphiphysin/Rvs (BAR) domain families respond to trafficking signals and dynamically transition between cytosolic localization and membrane-bound states ^13,14^. Once membrane bound, these proteins function in tubular-vesicular carrier formation and membrane fission ^15–21^. In many cases, membrane localization involves a short amino-acid sequence that changes from an unstructured state to an amphipathic helix ^22–24^. The amphiphysin N-BAR proteins, named for an N-terminal amino-acid sequence that can fold into an amphipathic alpha helix (called the H_0_ helix), are required for endocytic recycling and use a similar mechanism to transition from the cytosol to the membrane of the recycling endosome, where they promote membrane tubulation and fission during recycling carrier formation ^13,17,25,26^.

Progress has been made toward understanding the mechanism of recycling carrier formation, using *C. elegans* as a model system ^17,25,27,28^. Among the factors required for recycling carrier formation is AMPH-1, the sole amphiphysin protein in *C. elegans* ^25^. Like other amphiphysin orthologs, AMPH-1 is a dimeric, arc-shaped protein that previous studies have shown can induce membrane curvature and tubulation of negatively charged liposomes ^13,17,25,26^. These observations suggested a direct role for AMPH-1 in the creation of tubular-vesicular carriers. Recently, we demonstrated that not only can AMPH-1 induce membrane tubulation and fission *in vitro,* but that this activity is dramatically enhanced by guanine nucleotides ^17^. However, the mechanism underlying this nucleotide-stimulated membrane fission remains unresolved.

A number of physical models have been proposed to explain how peripheral membrane protein binding can induce bilayer deformation and fission, including amphipathic helix insertion and protein scaffold formation ^12,29,30^. Amphipathic helix insertion has been shown to induce membrane curvature, where the hydrophobic face of the helix creates a wedge in one leaflet of the membrane bilayer, altering its surface area relative to the other leaflet ^20,22,31,32^. In this wedge mechanism, amphipathic helix insertion impacts local curvature based, in part, on membrane-insertion depth: deep insertion results in negative curvature that can inhibit vesiculation, while shallow helix insertion induces positive curvature that is needed for tubulation and fission ^32–34^. Scaffold formation, by contrast, involves the oligomeric assembly of peripheral membrane proteins on the membrane surface in order to create a tubular structure via interactions between neighboring, membrane-bound protein units ^35–40^. Importantly, scaffold-induced tubulation has been proposed to create a diffusive restriction for lipids incorporated into the tubule, compared to those outside of it ^41,42^. The resulting diffusive inhomogeneity creates a frictional instability that accumulates tension as the tubule elongates, with the help of cytoskeletal proteins, until fission occurs ^43,44^.

Here, we show that GTP binding and hydrolysis alter the position of the AMPH-1 H_0_ helix, inducing a conformational change that regulates the membrane binding and tubulation necessary for fission. In free solution, AMPH-1 in the post-hydrolysis state (GDP), forms oligomeric assemblies via interaction between the H_0_ helices of separate homodimers. When AMPH-1 is trapped in the GTP-bound state (GMPPNP), oligomer formation in free solution is inhibited. At the same time, the GMPPNP-bound AMPH-1 readily binds membranes but is incapable of inducing tubulation. Remarkably, GTP hydrolysis results in formation of narrow membrane tubules that are stabilized by stacked AMPH-1 rings. We show that not only does nucleotide state influence the proximity of the H_0_ helices within and between homodimers in solution, but that nucleotide state affects membrane tubulation and scaffold formation, by altering both H_0_ helix position and depth of insertion in the membrane. These findings lead us to a model that involves aspects of both the wedge and scaffold models: GTP binding stabilizes a membrane-bound AMPH-1 homodimer with deeply embedded H_0_ helices, which promotes membrane binding but prevents premature tubule formation. GTP hydrolysis releases the H_0_ helix, resulting in more shallow membrane insertion, induction of positive membrane curvature and reorganization of the AMPH-1 dimers into a scaffold of stacked rings that are organized through interactions between H_0_ helices of separate AMPH-1 homodimers.

## Results

### Guanine nucleotide state links H_0_-helix interactions and AMPH-1 oligomer formation

The connection between guanine nucleotide state and changes in the conformation of the H_0_ helix is supported by our solution studies of the AMPH-1 deletion mutant (ΔH_0_), which is missing 16, N-terminal amino acids (Figure 1A). While thermostability measurements indicated that truncation does not impact structural stability (Figure S1A), ΔH_0_ failed to bind to liposome membranes in a co-sedimentation assay (Figure 1B; see also ^37^, or to form tubules, as assessed by negative-staining electron microscopy (Figure S1, C-E). ΔH_0_ displayed a significant decrease in nucleotide binding compared to AMPH-1, as observed by a decrease in fluorescence enhancement when bound to the fluorescent GTP analog, MANT-GTP (Figure 1D and ref ^17^) Likewise, ΔH_0_ displayed a significant reduction in the initial rate of GTP hydrolysis compared to AMPH-1 (Figure 1E and ref ^17^), suggesting a role, direct or indirect, for the H_0_ helix in nucleotide binding and hydrolysis. Conversely, we found that guanine-nucleotide binding and hydrolysis appears to regulate the role of H_0_-helices in AMPH-1 oligomer formation (Figure 2).

**Figure 1.**
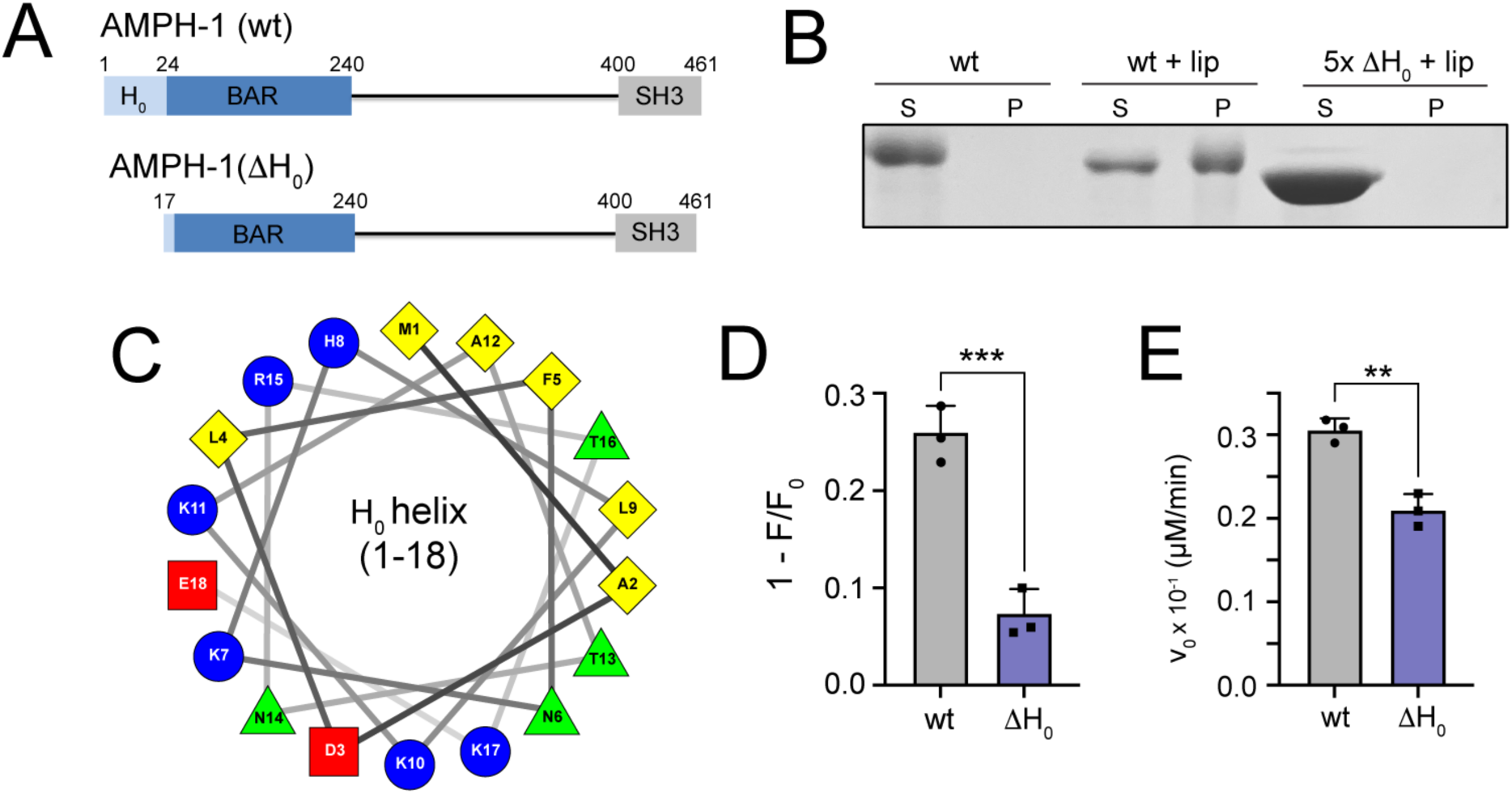
The H_0_ helix of AMPH-1 is important for both membrane and GTP binding. (A) Domain structures of full length wild-type AMPH-1 (wt) and a deletion mutant with the H_0_ helix removed (ΔH_0_) are shown. (B) Membrane binding by full length wt AMPH-1 and ΔH_0_ was assessed using a pelleting assay. Wt AMPH-1 (1 µM) and ΔH_0_ (5 µM) were mixed with 100% phosphatidylserine (PS) liposomes (200 nm; 0.33 mg/ml), incubated for 10 min and centrifuged at 90,000x g. Both supernatant (S) and pellet (P) fractions were recovered and examined by SDS-PAGE. (C) Helical wheel diagram of the AMPH-1 H_0_ helix. (D) Binding of GTP to wt AMPH-1 and ΔH_0_ was examined by fluorescence enhancement (1-F/F_0_) of 2 μM MANT-GTP in the presence of 50 µM of each protein ^17^. Bar plot shows the mean and error bars (± s.d.) of n = 3 independent experimental replicates. Significance was evaluated using a two-tailed Student’s t-test (***P = 0.001). (D) Steady state GTP hydrolysis (v_0_) by both full length and ΔH_0_ AMPH-1 (2.5 μM, in each case) was examined as previously described ^17^. Bar plot shows the mean and error bars (± s.d.) of n = 3 independent experimental replicates. Significance was evaluated using a two-tailed Student’s t-test (**P = 0.0023).

**Figure 2.**
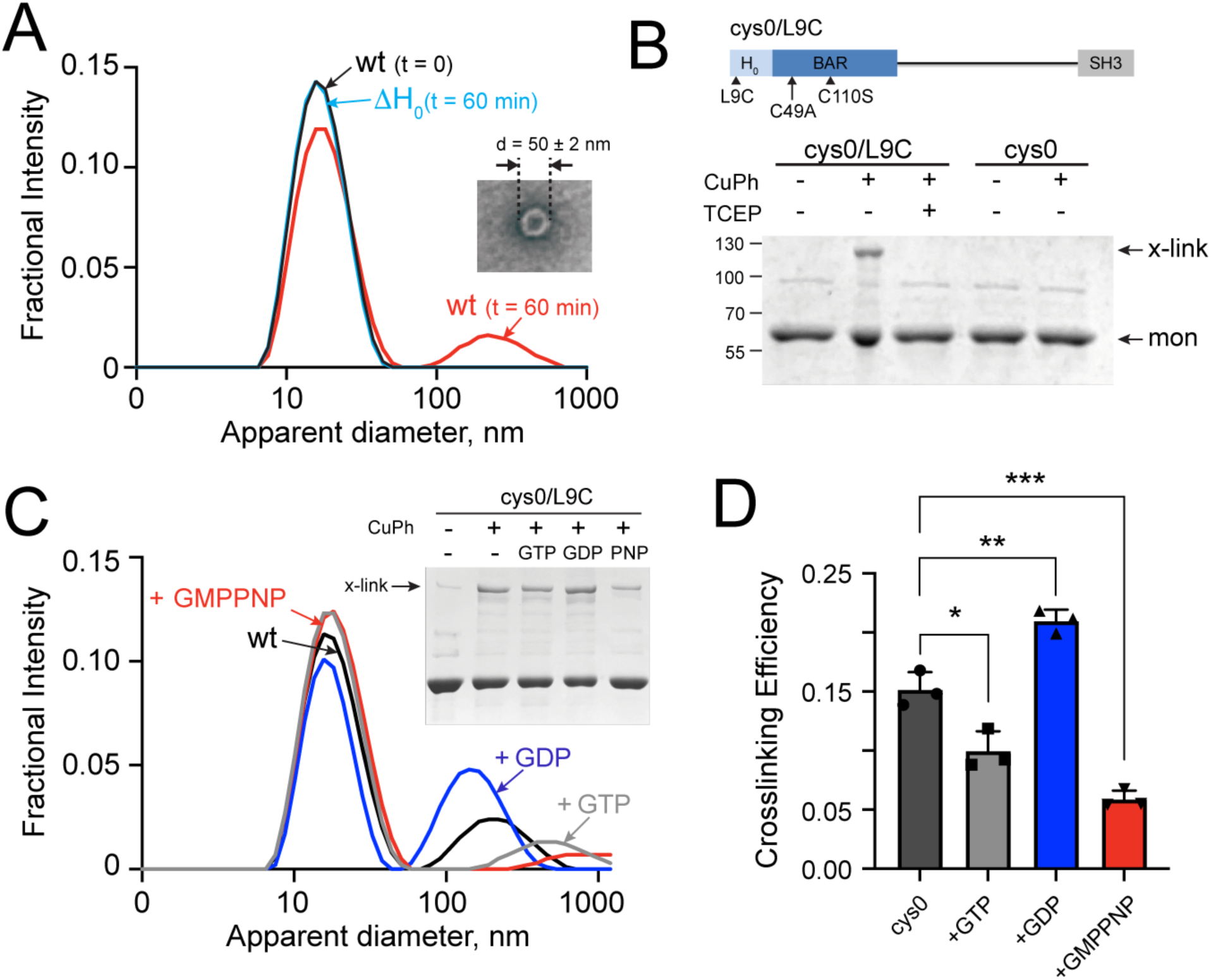
Formation of AMPH-1 oligomers in solution requires the H_0_ helix and is modulated by nucleotide state. (A) Apparent particle size distributions derived from dynamic light scattering (DLS) of wt AMPH-1 and ΔH_0_ are shown. In each case, samples (5 µM) were centrifuged (300,000x g for 30 min) to remove large multimers or aggregates and then examined by DLS either immediately or following a 60 min incubation at 23 °C. Particle size distribution for wt AMPH-1 both before (black) and after (red) a 60 min incubation are shown. The particle size distribution for ΔH_0_ after a 60 min incubation (light blue) is shown. The distribution for ΔH_0_ immediately after centrifugation (not shown) was identical to that observed after 60 min. The inset shows a negative stain EM image of an annular oligomer that is observed in higher molecular weight AMPH-1 complexes purified by gel filtration chromatography (see also Figure S2). (B) The domain structure of a cysteine free (cys0) variant of AMPH-1 containing a single cysteine at position 9 (L9C) in the H_0_ helix (cys0/L9C) is shown. Following centrifugation (300,000x g for 30 min), samples of either cys0/L9C or cys0 (2 µM, in each case) were incubated in the presence or absence of the strong thiol oxidant copper phenanthroline (CuPh, 1 mM) for 10 min at 23 °C. Samples were then treated with either EDTA (10 mM) alone or EDTA plus TCEP (1 mM) and examined by non-reducing SDS-PAGE. Migration positions of monomer (mon) and disulfide cross-linked dimer (x-link) are indicated. (C) Apparent particle size distributions derived from DLS of wt AMPH-1 in the presence of different guanine nucleotides. Following centrifugation, wt AMPH-1 (5 µM) was incubated either in the absence (wt; black) or presence of 500 µM GTP (gray), GDP (blue), or GMPPNP (red) for 60 minutes at 23 °C. The inset shows a disulfide cross-linking experiment using cys0/L9C AMPH-1 in the presence or absence of different guanine nucleotides. Sample preparation, centrifugation, oxidation and SDS-PAGE analysis were identical to panel (B). In each case, 2 µM protein was incubated with the indicated nucleotide (500 µM) for 60 minutes at 23 °C prior to oxidation and SDS-PAGE analysis. (D) The observed crosslinking efficiency of wt-cys0/L9C in the presence and absence of guanine nucleotides was quantified by densitometry (calculated as cross-linked dimer intensity / total intensity). The plot shows the observed mean and error bars (± s.d) of n = 3 independent experimental replicates like that shown in the panel (C) inset. Significance was assessed with a one-way ANOVA using the Geyser-Greenhouse correction (*P = 0.0157, **P = 0.0042, ***P = 0.0006).

To observe oligomer formation in solution, we used analytical, size-exclusion chromatography (SEC). SEC revealed two, well-separated peaks: an earlier peak, containing one or more oligomeric states of AMPH-1, and a later peak at the expected elution time for the AMPH-1 homodimer (Figure S2A). To examine the morphology of the oligomeric state(s), we isolated the earlier peak and performed negative-stain electron microscopy. The results suggest a range of assembled states, including both aggregates and distinct, ring-like structures (Figure 2A, inset; Figure S2B). To measure the particle size distribution of AMPH-1 under different conditions, we used Dynamic Light Scattering (DLS). Following removal of oligomers by ultracentrifugation (Figure S2C), AMPH-1 homodimers were incubated at room temperature for 60 min, and oligomer formation was monitored by changes in scattering profile. A time-dependent formation of large AMPH-1 complexes was observed (Figure 2A). To assess the role of the H_0_ helix in oligomer formation, similar experiments were conducted with ΔH_0_. No large complexes were observed by DLS for ΔH_0_, even after a 60 min incubation at 23°C. (Figure 2A). Moreover, SEC analysis demonstrated that ΔH_0_ eluted as a dimeric species (Figure S2A), and an AMPH-1 mutant, L9Q (Figure S2A), which displays reduced membrane fission activity ^17^, had a reduced oligomeric peak, as expected if the amphipathicity of the N-terminal H_0_ helix of AMPH-1 is required for oligomer formation (Figure S2A).

To determine whether oligomer formation in free solution involves H_0_ helix interactions between homodimers, we performed a disulfide cross linking experiment. A cysteine residue was introduced at position L9 of a cysteine-free of AMPH-1 (AMPH-1 C49A/C110S; referred to as “cys0”; Figure 2B), cys0/L9C. To promote the formation of disulfide bonds, purified cys0/L9C was incubated at room temperature for 60 min, then exposed to the oxidizing agent, copper phenanthroline (CuPh) for 5 minutes. Non-reducing SDS-PAGE analysis revealed a band the molecular weight of an AMPH-1 dimer that was susceptible to reduction by TCEP and absent in cys0 (Figure 2B). Importantly, the distance between H_0_ helices within a homodimer precludes intradimer crosslinking; therefore, the cross-linked band can only result from an interaction between H_0_ helices of separate AMPH-1 homodimers.

To examine the role of guanine nucleotides in oligomer formation in solution, we looked for increased particle size by dynamic light scattering (DLS) and nucleotide-dependent H_0_ helix interactions, using disulfide crosslinking. We found that guanine-nucleotide regulation of AMPH-1 oligomer formation in solution requires the H_0_ helices: the GTP-bound state (formed with GMPPNP) prevented interactions between AMPH-1 homodimers (Figure 2C), while GDP promoted ring-like oligomer formation in the presence (Figure 2A, inset), but not in the absence, of the H_0_ helices (Figure S2D). If guanine nucleotide-state regulates interactions between H_0_ helices of separate AMPH-1 homodimers, we would expect disulfide-bond formation by CuPh oxidation of H_0_ helix mutant, L9C (Figure 2B), to increase with GDP and decrease with GMPPNP, as observed (Figure 2C, D).

The impact of guanine nucleotides on both DLS and disulfide cross-linking experiments suggested that nucleotide state modulates the position of the H_0_ helix. To test this hypothesis, we examined the mobility of the H_0_ helix in solution using fluorescence anisotropy. We used a thiol alkylating fluorescent probe, IAEDANS, to covalently label position 11 (cys0/K11C; Figure 3A) on the hydrophilic face of the amphipathic H_0_ helix. The low, steady-state fluorescence anisotropy (0.04-0.07) observed, indicative of high mobility of the IAEDANS probe, is consistent with previous observations that the H_0_ helix of N-BAR proteins is poorly structured in solution, only becoming helical upon membrane binding ^21^. Notably, addition of GDP results in an increase in anisotropy, suggesting that the H_0_ helix becomes less mobile in the GDP-bound state of cys0/K11C (Figure 3A), even in solution.

**Figure 3.**
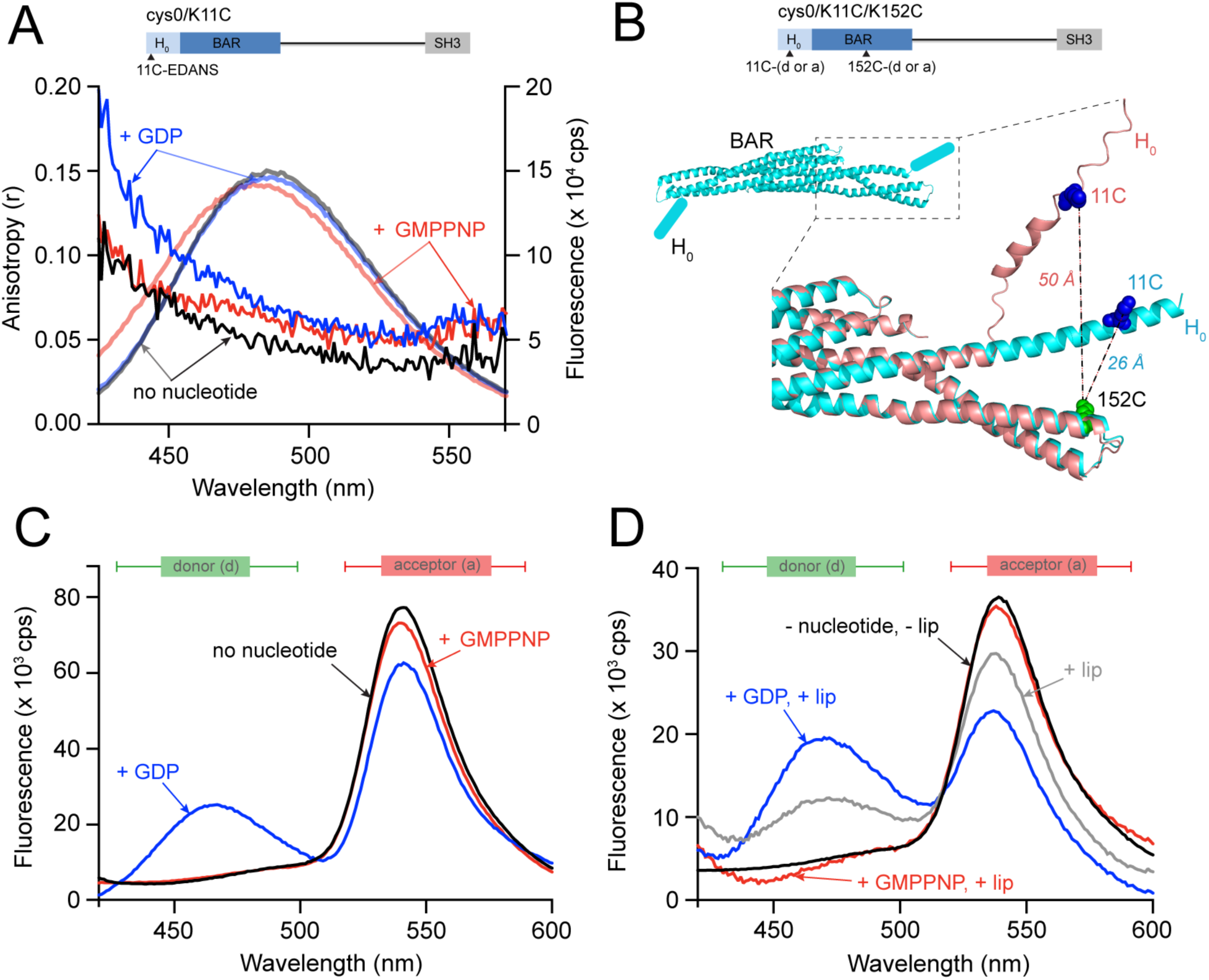
The dynamics and position of the H_0_ helix depends on nucleotide state. (A) Domain structure of AMPH-1 cys0 with a single Cys incorporated into the H_0_ helix (K11C) that was labeled with the fluorescent dye IAEDANS (cys0/11C-EDANS). The total fluorescence intensity and steady state anisotropy (lambda ex = 336 ± 2 nm) of cys0/11C-EDANS (0.1 µM) was examined in the absence of nucleotide (r, black; F_tot_, gray), or following addition of 1 mM GDP (r, dark blue; F_tot_, light blue) or 1 mM GMPPNP (r, dark red; F_tot_, light red). (B) Domain structure of AMPH-1 cys0 showing the incorporation positions of two Cys residues (K11C) and (K152C) used to attach the fluorescent dyes EDANS (donor; d) and fluorescein-5-maleimide (acceptor; a), which form a FRET pair when the dyes are within ∼ 25-65 Å of one another. The crystal structure of human amphiphysin (isoform 1) BAR domain dimer (inset; cyan; PDBID: 4ATM), viewed from below the membrane binding surface, illustrates the approximate location of the H_0_ helix (cylinder) at the tips of the BAR domain dimer. The relative positions and separations of the K11C and K152C mutations in an overlay of two, high-rank AlphaFold2 (Jumper et al., 2021) AMPH-1 structures (cyan and salmon) are shown in close-up. (C) The fluorescence emission spectra (lambda ex = 336 ± 2 nm) of double-labeled cys0/K11C/K152C (0.1 µM) mixed with 10 µM unlabeled AMPH-1 in free solution, in the absence of nucleotides (black), or following addition of 1 mM GDP (blue) or GMPPNP (red), is shown. (D) Fluorescence emission spectra of double-labeled cys0/K11C/K152C (0.1 µM) mixed with 10 µM unlabeled AMPH-1 in the presence of 200 nm, 100 % PS liposomes, with and without added nucleotides. In free-solution, the donor emission from the double-labeled AMPH-1 is almost fully quenched (black), indicating a close average proximity of the labeled sites. Addition of PS liposomes (gray) results in partial dequenching of the donor and reduction in sensitized emission from the acceptor of the double-labeled cys0/K11C/K152C AMPH-1, which increases in magnitude when 500 µM GDP is also present (blue). In the presence of both PS liposomes and 500 µM GMPPNP (red), little change in the relative emission of both donor and acceptor is observed, with the donor as fully quenched as in free solution.

To examine nucleotide-dependent changes in H_0_ helix dynamics within the AMPH-1 homodimer, we monitored the proximity between the H_0_ helix and the BAR domain, using Förster resonance energy transfer (FRET). To label AMPH-1 with thiol-reactive fluorescent donor and acceptor molecules, we introduced cysteine residues into cys0 at positions K11 (H_0_ helix) and K152 (BAR domain), generating AMPH-1 variant, cys0/K11C/K152C (Figure 3B), which was then labeled with IAEDANS (donor) and fluorescein (acceptor; Förster radius ∼46 Å). To minimize inter-dimer FRET, 10 µM AMPH-1 (unlabeled) was mixed with 100 nM of dual-labeled cys0/K11C/K152C. In the absence of nucleotides, the donor fluorescence is almost completely quenched, while the acceptor fluorescence is enhanced, relative to reference samples treated with protease to fully separate the dyes (Figure S3). This result demonstrates that the two probes are, on average, in close proximity to one another when AMPH-1 is in free solution (Figure 3C). Strikingly, while the presence of GMPPNP results in an average proximity between the two labeled positions that is similar to that observed in the absence of nucleotide, the addition of GDP resulted in a large dequenching of the donor and loss of sensitized fluorescence from the acceptor, indicating that GDP induces a significant increase in the average distance between the labeled positions on the H_0_ helix and BAR domain (Figure 3C).

### Membrane scaffold formation depends on guanine nucleotide state and H_0_ helix position

Like other N-BAR proteins, AMPH-1 homodimers require the H_0_ helices for membrane binding ^37,45^, yet when bound to membranes in the GTP-bound state (Figure S1B, AMPH-1 plus GMPPNP) membrane tubulation is inhibited ^17^. This observation, in combination with the nucleotide modulation of both oligomer assembly (Figure 2) and H_0_ helix dynamics in free solution (Figure 3), suggested that nucleotide hydrolysis is required to change the position of the H_0_ helices in favor of membrane tubulation. To test this hypothesis, we first examined the overall conformation of membrane-bound AMPH-1 using limited proteolysis in the absence and presence of 200 nm PS liposomes and different nucleotides. SDS-PAGE revealed distinct proteolytic AMPH-1 fragments, including a larger fragment in the presence of liposomes that is consistent with protection of the H_0_ helix from proteolysis (Figure S4A). Furthermore, we observed that addition of GMPPNP resulted in more protection from proteolysis, compared to proteolysis in the absence or presence of other guanine nucleotides (Figure S4B).

To examine guanine nucleotide-dependent changes consistent with formation of an oligomeric, AMPH-1 scaffold for tubulation, we again employed FRET with the dual-labeled cys0/K11C/K152C variant. For these experiments, 100 nM dual-labeled cys0/K11C/K152C was incubated with 10 µM unlabeled AMPH-1 (to minimize inter-dimer FRET) and 200 nm PS at room temperature, with or without guanine nucleotides. In the absence of nucleotide, the observed FRET signal displayed a small decrease (compared to the protein in free solution), suggesting that membrane binding results in a modest increase in the average distance between the donor and acceptor probes within the homodimer (Figure 3D). By contrast, in the presence of GMPPNP, the observed FRET signal is almost identical to that seen in free solution with GMPPNP, indicating that the donor and acceptor probes are in close proximity. Strikingly, addition of GDP causes a dramatic reduction in FRET efficiency, indicating a substantial increase in the inter-probe distance for membrane-bound AMPH-1 in the GDP-bound state, compared to nucleotide-free, or GTP-bound (GMPPNP) conformations.

Studies of the N-BAR protein, endophilin, suggested that direct physical contact between H_0_ helices plays a role in protein lattice assembly and the associated induction of membrane curvature ^36,37^. We reasoned that coupling of the AMPH-1 H_0_ helices to nucleotide hydrolysis could provide a regulatory mechanism to control H_0_ helical contacts between AMPH-1 dimers at the membrane surface. To test this hypothesis, we introduced a cysteine residue to either the hydrophobic (L9C), or hydrophilic (K11C) face of the AMPH-1 cys0 H_0_ helix and performed crosslinking experiments in the presence of liposomes. Either cys0/L9C, or cys0/K11C, was incubated with or without guanine nucleotides in the presence of liposomes, followed by oxidation with CuPh (Figure 4A). Our results revealed that GDP increased the crosslinking efficiency for cys0/L9C (Figure 4B, D), but significantly decreased the crosslinking efficiency for cys0/K11C (Figure 4C, E). In keeping with our findings that GDP and GMPPNP induce different changes to the orientation of the H_0_ helix in solution, the effects of GMPPNP on cys0/L9C and cys0/K11C crosslinking efficiencies on membranes were opposite to those of GDP (Figure 4D, E). To test the importance of H_0_-helix amphipathicity on nucleotide-dependent H_0_ helix interactions, we used the membrane-fission mutant, L9Q ^17^, with a cystine introduced into the H_0_ helix at position F5. Regardless of nucleotide state, the crosslinking efficiency for cys0/L9Q/F5C on membranes was significantly decreased, indicating that, with disruption of amphipathicity, GDP and GMPPNP could no longer regulate H_0_ helix interactions (Figure 4F, G). These crosslinking results support FRET experiments and are consistent with the H_0_ helices breaking away from the AMPH-1 BAR domain in the GDP-bound conformation (Figure 3), where they are available to form interactions with H_0_ helices of other AMPH-1 homodimers.

**Figure 4.**
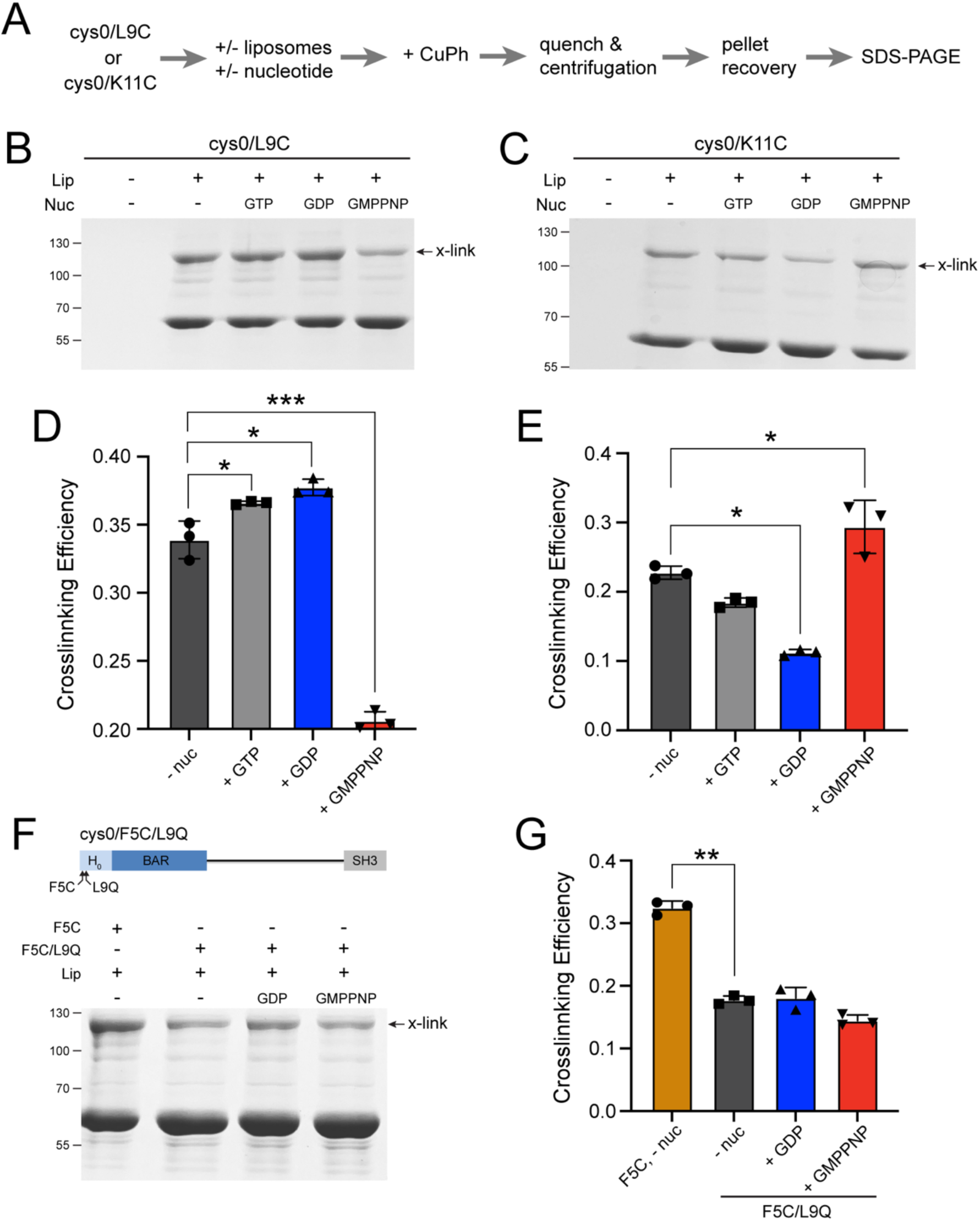
Nucleotide modulates the proximity and orientation of H_0_ helices at the membrane surface. (A) Schematic illustrating a disulfide crosslinking assay for probing the local H_0_ helix proximity of AMPH-1 at the membrane surface. (B) Disulfide cross linking was performed on wt-cys0/L9C in the presence of 200 nm, 100% PS liposomes and different guanine nucleotides. Following centrifugation (300,000x g for 30 min), the protein (10 μM) was incubated with 500 µM GTP, GDP, or GMPPNP in the presence of 0.33 mg/mL PS liposomes for 15 minutes at 23 °C. Each sample was then treated with 1 mM CuPh for 5 min, quenched with 10 mM EDTA and centrifuged (20,000x g) for 20 minutes at 23°C to recover the membrane-bound fraction. Pellets were recovered and analyzed using non-reducing SDS-PAGE. The cross-linked dimer species (x-link) is indicated. (C) The same protocol and reagent concentrations were used to examine the disulfide cross linking of membrane-bound wt-cys0/K11C. The crosslinking efficiency of (D) cys0/L9C and (E) cys0/K11C was determined using densitometry and calculated as the ratio of the cross-linked dimer intensity to the sum of the total AMPH-1 intensity (x-linked dimer plus monomer). Plots shows the mean and error bars (± s.d) of n = 3 independent experimental replicates. Significance was assessed with a one-way ANOVA using the Geyser-Greenhouse correction. For panel (D), *P = 0.0281, *P = 0.0115, ***P = 0.00094. For panel (E), *P = 0.023, *P = 0.0444. (F) Domain structure of AMPH-1 cys0 incorporating both a single Cys in the H_0_ helix (F5C; cys0/F5C) and a membrane binding and fission inhibiting mutation ^17^, L9Q (cys0/F5C/L9Q). The same protocol and reagent concentrations were used to examine the disulfide cross linking of membrane-bound cys0/F5C and cys0/F5C/L9Q. (G) Quantification of crosslinking efficiency for cys0/F5C and cys0/F5C/L9Q. The plot shows the mean and error bars (± s.d) of n = 3 independent experimental replicates. Significance was assessed with a one-way ANOVA using the Geyser-Greenhouse correction with **P = 0.0034.

### Guanine nucleotide state alters H_0_ helix membrane penetration and membrane curvature

To test for membrane insertion of the H_0_ helix, we exploited a well-established fluorescence membrane penetration assay based on the environmentally sensitive fluorescent dye NBD ^46–48^. In polar environments, like aqueous solution, the NBD probe is quenched, while in apolar or hydrophobic environments, the dye becomes highly fluorescent ^47,49^. We labeled AMPH-1 with an iodo-acetamido version of NBD (IANBD) on the hydrophobic H_0_-helix position, F5, using the cys0/F5C variant (Figure 5A). When compared to the fluorescence emission of NBD labeled cys0/F5C in solution, in the presence of liposomes, the fluorescence emission shifted to shorter wavelengths and increased in intensity (Figure 5C), consistent with a transition of the NBD probe from an aqueous to a hydrophobic environment. Addition of GMPPNP resulted in an approximately 6-fold fluorescent enhancement, while GDP resulted in an approximately 2-fold reduction in fluorescence (Figure 5D). Similar results are obtained when the composition of the liposomes is changed to a mixture of PS and PC (Figure S5A). To eliminate the possibility that a hydrophobic patch on the protein caused the changes in NBD fluorescence, we repeated the experiment with a nitroxide-labeled, fluorescence quencher in the membrane. The nitroxide-label, 16:0-5 doxyl-PC, was incorporated into liposomes (Figure 5B) and mixed with NBD-labeled cys0/F5C and nucleotide. Upon incubation with GMPPNP, NBD fluorescence was significantly quenched, whereas GDP did not result in a significant change in NBD fluorescence (Figure 5E, F), suggesting that the H_0_ helix of AMPH-1 is more deeply inserted into the membrane in the GTP-bound state than in the GDP-bound state.

**Figure 5.**
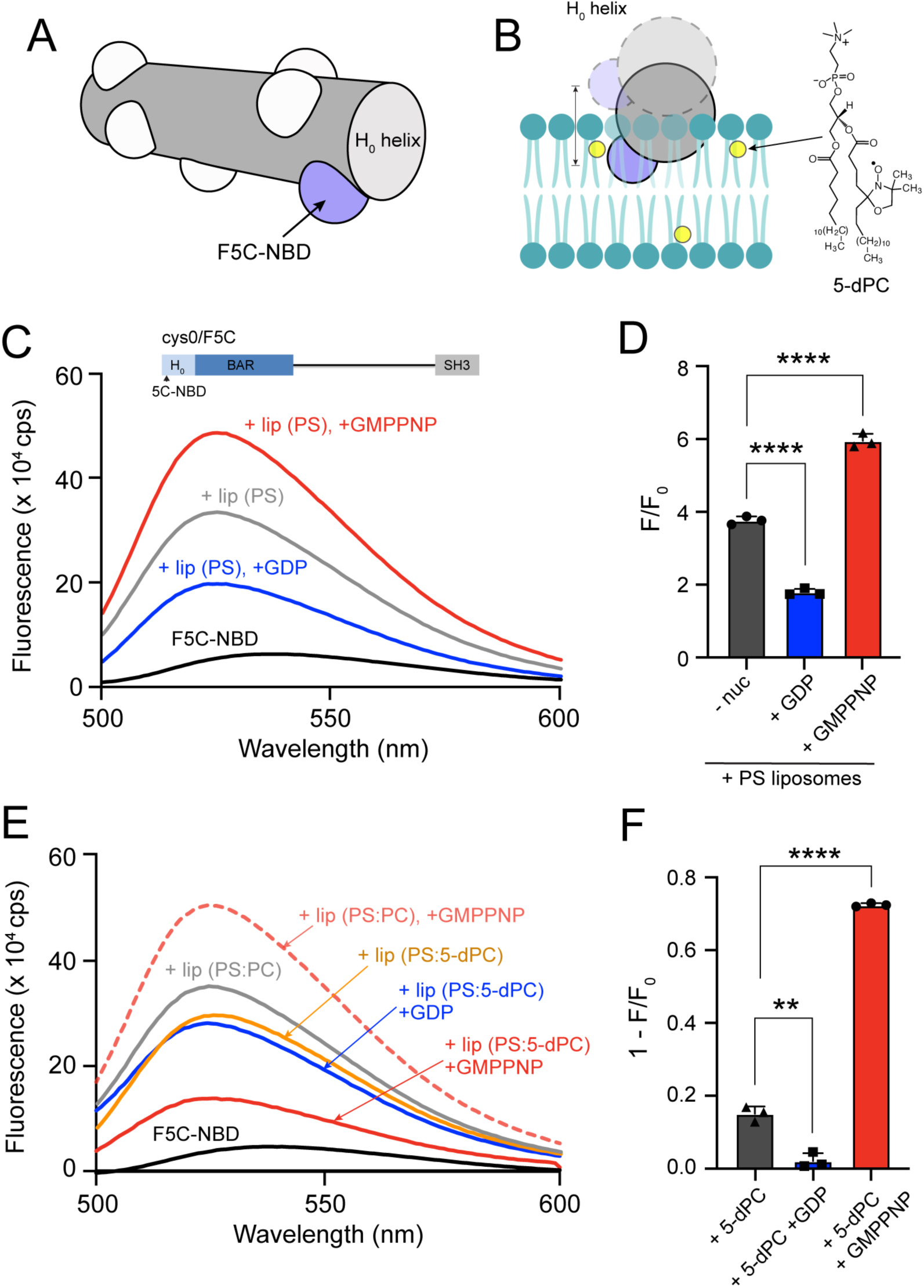
GDP stabilizes a shallower penetration of the H_0_ helix into the lipid bilayer. (A) Model of the AMPH-1 H_0_ helix showing the attachment position (F5C) of the fluorescent dye IANBD (NBD). (B) The average proximity of the NBD probe and a nitroxide quencher, which is incorporated into the C5 position of one fatty acid side chain of phosphatidylcholine (5-dPC), is predicted to change if the average bilayer penetration of the H_0_ hydrophobic face is altered by nucleotide state. (C) Steady state fluorescence emission spectra (lambda ex = 480 ± 2 nm) for cys0/F5C-NBD (0.1 µM) in the absence (black) and presence of 200 nm, 100% PS liposomes (PS; 0.3mg/mL; gray). Addition of GDP (500 µM; blue) results in increased quenching of the NBD probe, consistent with a reduction in average helical penetration into the membrane. The presence of GMPPNP (500 µM, red) results in a large increase in fluorescence, consistent with an increase in average helical penetration. (C) Quantitation of NBD fluorescence enhancement (F/F_0_) with plot showing the mean and error bars (± s.d) of n = 3 independent experimental replicates. Significance was assessed with a one-way ANOVA using the Geyser-Greenhouse correction with ****P<0.0001. (D) Molecular structure of fluorescence quencher-modified PC, 5-doxyl-phosphatidylcholine (5-dPC), used to examine membrane penetration depth of the NBD probe. Steady state fluorescence emission spectra (lambda ex = 480 ± 2 nm) for cys0/F5C-NBD (0.1 µM) in the absence of liposomes (black) and in the presence of PS:PC (0.9:0.1) or PS:5-dPC (0.9:0.1) liposomes (0.3 mg/ml), with or without 500 nM GDP or 500 nM GMPPNP. The fluorescence of the NBD probe upon addition of PC:PS liposomes only (gray), PC:5dPC liposomes only (orange), PC:5dPC liposomes with GDP (blue) and PC:5dPC liposomes with GMPPNP (red) are shown, with the fluorescence enhancement of the NBD probe in the presence of PS:PC liposomes and 500 nM GMPPNP (dashed red) shown for reference. (E) Quantitation of 5-dPC-induced quenching for cys0/F5C-NBD for PS:5-dPC liposomes, relative to the emission for PS:PC liposomes under identical conditions (1-F/F_0_), is shown as the mean and error bars (± s.d) of n = 3 independent experimental replicates. Significance was assessed with a one-way ANOVA using the Geyser-Greenhouse correction with ****P =0.0002 and **P = 0.0015.

In a complementary approach, we employed a different fluorescent probe (EDANS) as a sensor of H_0_ helix insertion into the membrane. In this case, the expected fluorescence response is opposite to that observed with NBD, with the EDANS probe displaying reduced fluorescence upon insertion into a lipid bilayer ^50^. In the presence of PS liposomes, the EDANS-labeled cys0/F5C AMPH-1 variant displayed an approximately 13% reduction in fluorescence compared to the labeled protein in free solution (Figure S5A). Addition of GMPPNP resulted in an increase in observed fluorescence quenching (approximately 20% total reduction, compared to the protein in free solution; Figure S5B and C), as expected if the H_0_ helix is inserted more deeply into the membrane. By contrast, addition of GDP resulted in a reduced quenching of the EDANS probe (approximately 6% total reduction, compared to the protein in free solution; Figure S5B and C). The increased quenching observed in the presence of GMPPNP and reduced quenching seen with GDP are consistent with nucleotide-directed movement of the H_0_ helix deeper into and further out of the membrane, respectively. To test the importance of H_0_-helix amphipathicity on nucleotide-dependent membrane insertion, we introduced the L9Q mutation into cys0/F5C, creating cys0/F5C/L9Q. When IAEDANS labeled cys0/F5C/L9Q was mixed with liposomes, we observed a reduced quenching of the probe in the presence of PS liposomes (approximately 6% reduction, compared to the protein in free solution; Figure S5D). Importantly, neither addition of GMPPNP, nor GDP, changed the observed level of fluorescence quenching, indicating that the L9Q mutation greatly reduced the sensitivity of membrane insertion to guanine nucleotides (Figure S5D).

### Structure of AMPH-1-induced tubules reveals an H_0_ helix-linked lattice of stacked rings

Previously, we observed by negative-stain electron microscopy rigid, tubular membrane structures formed by liposomes incubated with AMPH-1 and GTP ^17^ (Figure 6A). To visualize in greater detail how AMPH-1 forms scaffolds on membrane tubules in the presence of nucleotides, we examined these tubules using cryo-EM single particle reconstruction. AMPH-1 was incubated with large (1 µm) PS liposomes and GTP for 60 min prior to vitrification. Because our previous data showed that AMPH-1 cannot form tubules in the presence of the non-hydrolyzable GTP analog, GMPPNP, and because tubules formed in the absence of nucleotide are highly irregular and heterogeneous ^17^, we anticipated that the most homogenous tubular structures would likely represent the post-hydrolysis, GDP-bound, membrane-associated AMPH-1. Imaging revealed a large population of tubules, distributed over a range of lengths and widths, with a substantial sub-population displaying a narrow and relatively consistent geometry (Figure 6A). This sub-population was isolated, segmented and subjected to 2D classification analysis (Figure S6A), resulting in a total of 539,488 particles and 20 class averages (Figure S5B).

**Figure 6.**
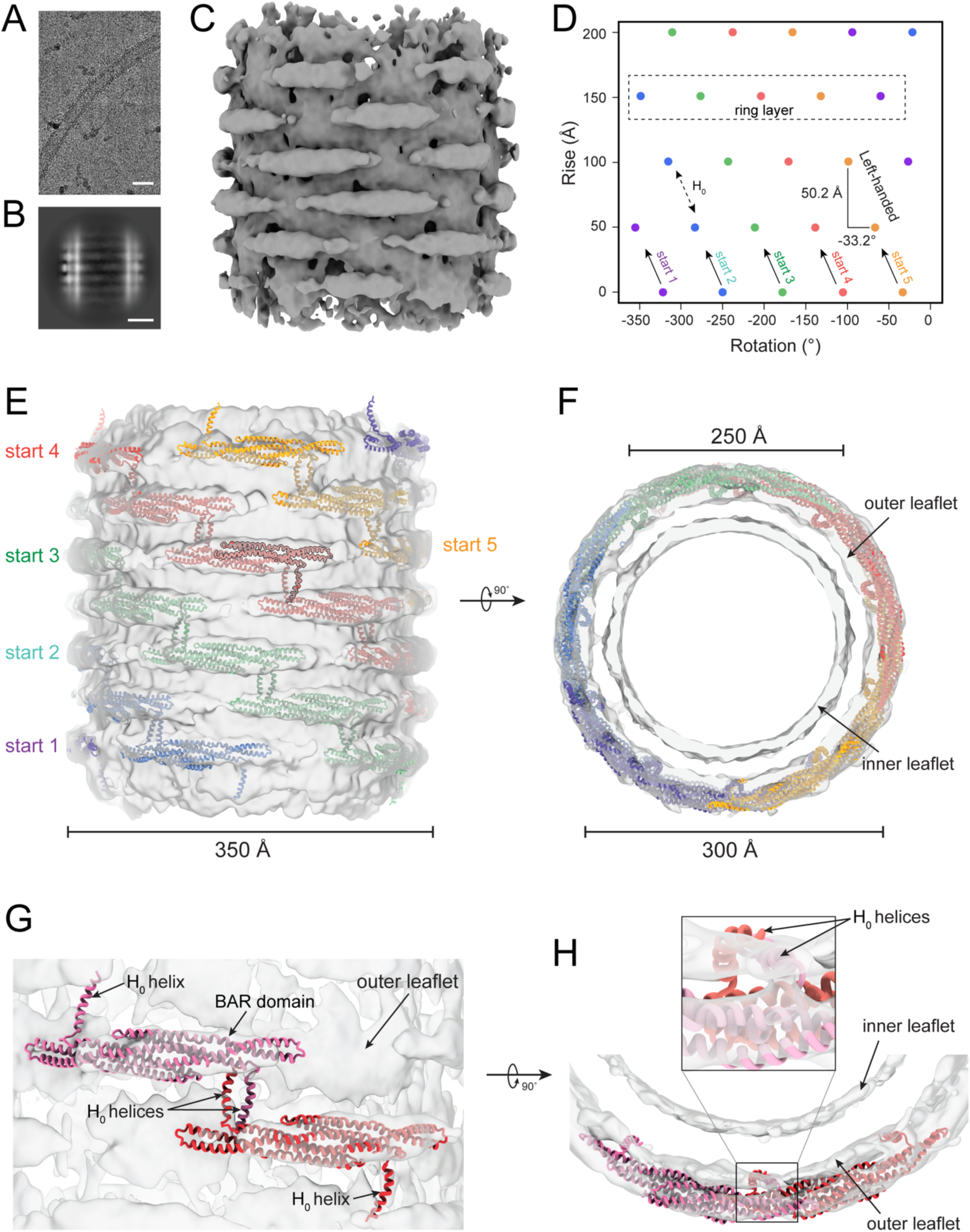
CryoEM structure of AMPH-1 membrane tubule formed after extended incubation with GTP. (A) A representative electron cryo-micrograph (motion corrected) of membrane tubules obtained by mixing AMPH-1 with 1 µm PS liposomes and GTP. Samples were vitrified following a 60 min incubation at 23 °C. Scale bar denotes 500 Å. (B) Selected 2D class average of the tubule showing the separation (50 Å) between each AMPH-1 layer. The scale bar denotes 500 Å. (C) Low-pass filtered cryo-EM density map obtained from the 2D class in (B), showing the well-resolved layers of AMPH-1 dimers. (D) Assembly pattern map showing the helical rise (50.2 Å) and twist angle (-33.2°) of the AMPH-1 dimers along the length of the tubule. Each dot represents a single subunit with the color keyed to the fitted model below. (E) Side view of the cryo-EM map fitted with a Alphafold2 predicted atomic model ^51^ of the AMPH-1 dimer, excluding the middle unstructured and C-terminal SH3 domains, which are not resolved in the map. Dimers are arranged in 5-membered stacked rings, with the rings rotationally offset (-33.2°) with respect to one another, leading to a helical pattern of dimers along the tubule length (purple, cyan, green, red, and orange). For reference, one AMPH-1 monomer in the third ring from the top (red) is outlined in black. The outer diameter of the AMPH-1-bound tubule is approximately 350 Å. (F) Top end-on view of the tubule showing that the diameter of the membrane tubule lumen (i.e., distance between the inner membrane leaflets) is approximately 250 Å, while the distance across the tubule between the outer leaflets is approximately 300 Å. (G) Close up view of two AMPH-1 dimers and predicted inter-dimer interaction via their H0 helices. The H_0_ helices of each dimer project orthogonally from the BAR domains and point in opposite directions along the long axis of the tubule. (H) Side view of the predicted H_0_ interaction region, showing the position of helices with respect to the outer leaflet of the lipid bilayer. Inset shows a closer view of the H_0_ helices from this orientation.

The overall AMPH-1 tubule morphology is consistent across the 2D class averages, though a range of tubule diameters and feature heterogeneities were observed. The most homogeneous class was selected for refinement (Figure 6B; Figure S6A) and subsequent single particle reconstruction yielded an approximately 8 Å map of a membrane tubule, wrapped in well-resolved rings of protein density (Figure 6C). The apparent twist-rise parameter pair (50.2 Å rise, -33.2° twist) was established using iterative helical refinement until a satisfactory map was obtained. The observed protein densities are highly consistent with the size and shape of the AMPH-1 dimer and are well fit by a predicted AlphaFold2 model ^51^ of the AMPH-1 N-BAR dimer (Figure 6E). Because the unstructured middle domain and C-terminal SH3 domain are not resolved in the map, these sequences were not included in the fitting. The protein-wrapped tubule is approximately 35 nm in external diameter and contains an internal membrane tubule, approximately 25 nm in diameter, with well resolved inner and outer lipid leaflets (Figure 6D). Because both right- and left-handed helical symmetries fit the data well, it was not possible to establish the handedness of the AMPH-1 lattice. Importantly, the relative positions of the BAR domains and H_0_ helices are almost identical for the two helical symmetries. The left-handed model is shown in Figure 6.

A striking feature of the pseudo-atomic model of the AMPH-1 tubule is the predicted location of the H_0_ helices. While the refined cryo-EM map cannot unambiguously identify the position of the H_0_ helices (likely because they cannot be distinguished from the lipid headgroup density at the membrane surface), fitting of a top-scoring AlphaFold model into the cryo-EM map results in localization of the H_0_ helices at the surface of the outer membrane leaflet, with the two helices of each BAR domain bent at nearly 70° angles from the BAR domain and pointing in opposite directions from one another (Figure 6E, G). At the same time, the H_0_ helices of each BAR domain appear to interact with the H_0_ helix of the rotationally offset BAR domains in the scaffold rings above and below (Figure 6D). This arrangement leads to the formation of five-membered, self-terminating AMPH-1 rings that stack directly atop one another. The localization of paired H_0_ helices at the membrane surface (Figure 6 G, H) in the presence of GDP is highly consistent with our biochemical characterization (Figure 4, 5).

## Discussion

Our previous findings suggest that AMPH-1 induces membrane fission through an unknown, guanine nucleotide-stimulated mechanism ^17^ that links GTP hydrolysis to AMPH-1 conformational changes, membrane tubulation, and fission. While other studies have implicated N-BAR H_0_ helices in membrane tubulation and fission ^22,31,32,36,52,53^, a role for guanine nucleotide binding and hydrolysis was unknown. For many ENTH and BAR domain proteins, the transition from cytosolic localization to a membrane-bound state involves rearrangement of a stretch of amino acids, which are unstructured in the cytosolic form, into an amphipathic helix upon membrane association ^20–24,54,55^. Here, we have established similar features for cytosolic and membrane-bound AMPH-1 and propose two additional states: (1) the transition from deep-to-shallow H_0_ helix membrane insertion upon GTP hydrolysis, and (2) interaction between H_0_ helices of neighboring homodimers in a scaffold-forming, GDP-bound set of stacked AMPH-1 rings.

Our FRET, crosslinking and cryoEM studies support nucleotide-dependent scaffold formation by AMPH-1. In the GTP-bound state, AMPH-1 scaffold formation is inhibited (Figure 3, 4), until hydrolysis has occurred, whereupon a scaffold forms through interactions between H_0_ helices of separate homodimers in the GDP-bound state (Figure 4, 6). At the same time, nucleotide hydrolysis impacts membrane curvature via changes in the depth of amphipathic helix insertion into the membrane. Based on our fluorescence quenching observations (Figures 5 and S5), the GTP-bound (GMPPNP) state of the AMPH-1 homodimer is trapped with the hydrophobic faces of its H_0_ helices inserted deeply into the membrane, an interaction that disfavors the positive curvature necessary for membrane tubule formation ^34,56^. By contrast, in the GDP-bound state, we observe shallow penetration of the H_0_ helices (Figure 5), which facilitates the positive curvature needed for tubule formation and fission ^31,34^.

Taken together, our data suggest a model in which highly curved, AMPH-1 scaffolds form on membrane surfaces in direct response to conformational changes that are linked to GTP binding and hydrolysis (Figure 7). In the GTP-bound state, AMPH-1 homodimers interact with negatively charged phospholipid bilayers through the positively charged face of the arc-shaped BAR domain ^57,58^ and the hydrophobic face of the amphipathic, N-terminal H_0_ helices, which are deeply inserted into the phospholipid bilayer (Figure 7C and E). We speculate that in this conformation, the deep penetration of the H_0_ helices induces negative curvature, which offsets any positive curvature favored by the BAR domain structure ^23,31,52^. By reducing or canceling the intrinsic positive curvature of the BAR domain, deep H_0_ helical insertion permits high local concentrations of AMPH-1 to accumulate at the membrane surface without prematurely creating the positive membrane curvature needed to drive cooperative scaffold assembly ^22,59,60^. Moreover, in the GTP-bound state, interactions between AMPH-1 homodimers are precluded by burial of the hydrophobic side chains of the amphipathic, H_0_ helices in the lipid bilayer.

**Figure 7.**
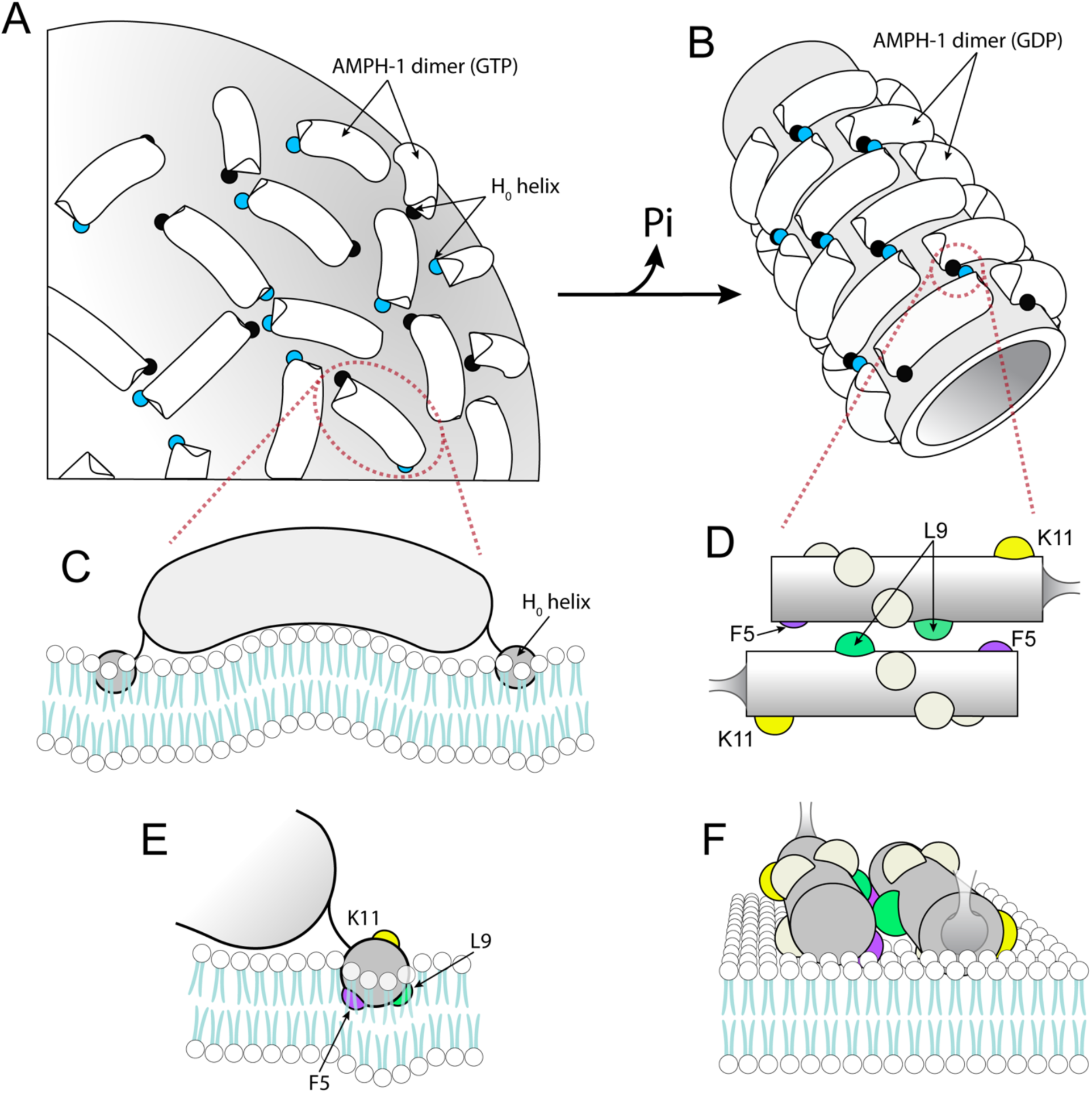
Model linking GTP hydrolysis to AMPH-1-induced membrane tubulation. (A) A model of AMPH-1 homodimers bound to the surface of electronegative phospholipid membranes in the GTP-bound state, which cannot form an organized lattice with persistent positive curvature. Homodimer BAR domains are shown as curved cylinders (white), with one N-terminal H_0_ helix of the dimer indicated as a blue and the other as a black circle. The position and dynamics of the unstructured middle and C-terminal SH3 domain are unresolved and are not shown. (B) Hydrolysis of GTP to GDP triggers reorganization of surface-bound AMPH-1 into stacked rings of 5 homodimers, which are connected to one another by hydrophobic re-alignment of the H_0_ helices. (C, E) GTP binding causes deeper penetration of the H_0_ helices into the membrane, preventing contact between the H_0_ helices of different AMPH-1 homodimers, blocking lattice formation. Deeper penetration may impose localized negative curvature ^32–34^ that offsets the positive curvature of the membrane binding surface of the BAR domain. (D, F) GTP hydrolysis causes a change in position and less membrane penetration, permitting the H_0_ helices of different AMPH-1 homodimers to interact at the bilayer surface (sideview, D; end view, F). Shallow penetration and helical contact act in concert with the curvature of the BAR domain surface to induce formation of stacked AMPH-1 rings and drive lattice formation, for narrow membrane tubules.

Following GTP hydrolysis, the H_0_ helices rotate to a shallower bilayer penetration depth (Figure 7B, D and F). In this position, the H_0_ helices favor the positive curvature needed to induce cooperative assembly of the BAR domain scaffold ^34,38,61,62^ and are freed to interact with one another through the same hydrophobic side chains that were buried in the bilayer prior to hydrolysis (Figure 7D, E and F; L9, yellow and F5, purple). Contacts between H_0_ helices both drive and stabilize formation of an oligomeric AMPH-1 lattice around the membrane tubule. In this GDP-bound state, the narrow tubule diameters (between 30-50 nm) observed by cryo EM are proposed to facilitate fusion of closely approximated, cross-tubule bilayers for membrane fission and carrier formation.

While membrane tubulation by N-BAR proteins has been observed previously ^18,21,22,31,63,64^, the lattice we observe for AMPH-1 following GTP hydrolysis is distinctive. Other N-BAR protein tubule structures, for example endophilin ^36^ and amphiphysin/BIN1 ^65^, display clear helical wrapping of the interacting BAR domains around the tubule exterior. The AMPH-1 lattice, however, is constructed from stacked rings of AMPH-1 dimers with no discernable helical pitch. This stacked-ring structure may be optimized to accommodate RME-1, a member of the EHD protein family and an AMPH-1 binding partner, known to be required for productive recycling carrier formation, *in vivo* ^25,27^. Tubules formed with both AMPH-1 and RME-1 are wider and have a lattice structure that is strikingly different from that formed by AMPH-1, alone ^25^. Recently, we observed that RME-1 dramatically slows AMPH-1 mediated membrane fission, *in vitro* ^17^ consistent with the finding that coordination between the two proteins is important for productive carrier formation during endocytic recycling, *in vivo* ^25^.

In light of our observations that membrane binding and tubulation by AMPH-1 is regulated by a guanine nucleotide hydrolytic cycle, it is interesting to note that GTPase-Activating Protein (GAP) and Guanine nucleotide Exchange Factor (GEF) domains have been identified in other BAR domain proteins ^13,66,67^. It remains to be determined how endocytic recycling is coupled to the nucleotide cycle of AMPH-1, and how carrier formation by fission of membrane tubules is regulated by other cellular factors, including EHD proteins like RME-1 ^17,25^ and cytoskeletal proteins ^68–70^. We expect future cryo-EM structures of the AMPH-1/RME-1 tubules trapped in various nucleotide states, plus high-resolution, *in vivo* images of the arrangement of these proteins on tubules of recycling endosomes, to shed light on this exquisitely regulated process.

## Methods

### Protein Mutagenesis, Expression and Purification

All mutations were performed using the Quick Change PCR protocol (Q5 Site-Directed Mutagenesis Kit E0554, NEB), and were based on an AMPH-1 pGEX-2T expression vector. *C. elegans* AMPH-1 constructs (wt, ΔH_0_, AMPH-1 L9Q, AMPH-1 C49A/C110S/F5C, AMPH-1 C49A/C110S/F5C/L9Q, AMPH-1 C49A/C110S/L9C, AMPH-1 C49A/C110S/K11C, AMPH-1 C49A/C110S/K11C/K152C) were expressed as N-terminal GST fusion proteins in *E. coli* Rosetta (DE3) cells using 0.4 mM isopropyl β-D-1-thiogalactopyranoside (IPTG) induction. Cell lysates were loaded onto a glutathione column equilibrated with buffer (50 mM Tris, pH 7.4, 300 mM NaCl, 1 mM MgCl2, 1 mM DTT). The eluted protein was cleaved with PreScission (GenScript) protease by dialysis and run over a second glutathione affinity column. The protein was then concentrated, flash-frozen, and stored at -80 °C. Protein oligomers and small aggregates that form during purification were removed by centrifugation at 300,000x g for in a TLA100 rotor using an Optima MAX-XP ultracentrifuge (Beckman) for 30 min at 4 °C prior to all biochemical experiments. Unless otherwise noted, all AMPH-1 concentrations listed in this manuscript are calculated for the dimer.

### Protein Labeling

Freshly purified AMPH-1 was diluted into preparation buffer (25 mM Tris pH 7.4, 300 mM NaCl, 1 mM MgCl_2_) and concentrated in a Vivaspin 4 MWCO 50,000 (Sartorius) at 4 °C to a final volume of 1-2 ml. The concentrated sample was diluted to a final volume of 15 ml in preparation buffer, concentrated and then diluted again for a total of up to three rounds of concentration and dilution, or until the initial sample was exchanged ∼500 to 1000-fold. The sample was then concentrated to a final volume of 2.5 ml and the buffer was exchanged using a PD-10 column equilibrated in reaction buffer (25 mM Tris pH 7.4, 150 mM KCl, 1 mM MgCl_2_) at room temperature. The final concentration of AMPH-1 was determined and the sample was transferred to a 5 ml conical reaction vial containing a spin vane on a magnetic stir plate. In all cases, aliquots of a reactive dye stock (≥ 10 mM in DMF) were added to AMPH-1 samples under low light conditions at 23 °C with stirring. For IAEDANS (Invitrogen/Molecular Probes), wt-cys0/K11C was labeled using two, sequential additions of the dye at a 3:1 dye:AMPH-1 monomer ratio (6:1 dye:monomer total mixing ratio), with incubations of 45 min in the dark at 23°C following each dye addition. For double labeling with IAEDANS and fluorescein, wt-cys0/K11C/K152C was prepared and first mixed with IAEDANS at a final dye:monomer ratio of 1:0.66. Following a 30 min incubation at 23 °C, fluorescein-5-maleimide (ThermoFisher/Invitrogen) was added at a 2:1 (dye:monomer) ratio and the reaction continued with stirring in the dark at 23 °C for an additional 40 min. Subsequently, IAEDANS and fluorescein were added at an additional 3:3:1 mixing ratio of IAEDANS:fluorescein:monomer and the same incubated for an additional 40 min at 23 °C in the dark. The protocol used for IANBD (Setareh Biotech) was identical to that used for IAEDANS labeling. In all cases, reactions were stopped by the addition of 5 mM reduced glutathione, followed by a 10 min incubation in the dark at 23 °C with stirring. Free dye removed using repeated dilution and concentration in a spin concentrator until an effective dilution of 2000 to 3000-fold was achieved, followed by final purification and buffer exchange using a PD-10 column. Labeled AMPH-1 samples were concentrated using a Vivaspin 4 MWCO 50,000 (Sartourius), snap frozen using liquid nitrogen and stored at −80 °C. Typical labeling efficiencies were ∼99% for IAEDANS, ∼98% for IANBD, and in the case of double labeling, ∼45% for IAEDANS and 49% for fluorescein.

### MANT-GTP Binding and GTPase Assays

GTP binding and turnover assays were conducted as previously described ^17^. Briefly, 50 µM AMPH-1 was incubated with 2 µM MANT-GTP (ThermoFisher/Invitrogen) prior to measurement in 25 mM Tris, pH 7.5, 100 mM KCl, 5 mM MgCl_2_. Emission spectra were acquired using a photon-counting T-format steady-state fluorometer with excitation at 356 nm and emission collected from 380 to 500 nm. All measurements were performed at 23 °C. GTP turnover was measured using a real-time inorganic phosphate assay (EnzCheck; ThermoFisher/Invitrogen) that is based on a colorimetric change of 2-amino-6-mercapto-7-methylpurine riboside following its phosphorylation by purine nucleoside phosphorylase ^71^.

### Dynamic Light Scattering

The hydrodynamic diameter of oligomers formed by wt AMPH-1 and ΔH_0_ in the presence of 500 µM guanine nucleotides was examined using a Zetasizer Nano (Malvern) DLS instrument (630 nm at a scattering angle of 173°). Prior to measurement, all samples were filtered through 0.2 μm membrane filters. DLS measurements were conducted on 100 μl samples of 5 µM protein in a BRAND UV cuvette. DLS curves were calculated from the average of 20, 120 s scans.

### Fluorescence Anisotropy

Fluorescence anisotropy measurements were conducted on AMPH-1 C49A/C110S/K11C, labeled with IAEDANS (100 nM), mixed with 10 µM unlabeled wt AMPH-1 in the presence or absence of GDP or GMPPNP. Excitation was performed using a wavelength of 337 nm with a 2 nm bandpass. For anisotropy measurements, each detection channel utilized the same emission filter set and a film polarizer.

### Liposome Preparation, Co-sedimentation and Tubulation Assays

Liposomes were prepared using a previously described protocol ^17,19^. Briefly, samples of brain extract phosphatidylserine (PS; Avanti) in solvent were first dried using a dry argon stream, followed by overnight incubation under vacuum at 23 °C. Dried lipids were resuspended in argon-sparged liposome extrusion buffer (50 mM HEPES pH 7.4, 150 mM NaCl, 25 mM KCl) to a final concentration of 1 mM followed by 10 successive freeze/thaw rounds using liquid nitrogen and a 52-60 °C water bath. Unilamellar liposomes were prepared by extrusion of the hydrated lipid mixture through Nuclepore track-etched polycarbonate filter disks (Whatman) using 11 rounds of successive processing in an Avanti Mini-extruder. The resulting liposome suspension was centrifuged at 20,000x g for 30 min at 21°C and the supernatant was discarded. Liposome pellets were then gently resuspended in liposome extrusion buffer. Lipid mixtures consisted of: (1) 100% brain extract phophosphatidylserine (PS; Avanti, cat. 840032); (2) 90% PS and 10% 1,2-dipalmitoyl-sn-glycero-3-phosphocholine (PC; Avanti, cat. 850355); (3) 90% PS and 10% 1-palmitoyl-2-stearoyl-(5-doxyl)-sn-glycero-3-phosphocholine (16:0-5 Doxyl PC; Avanti, cat. 810601).

For co-sedimentation assays, 2 µM protein was incubated with 0.4 mg/mL 200 nm liposomes in 100 µL of buffer in the presence or absent of 500 µM guanine nucleotides for 10 min prior to sedimentation at 90,000x g for 15 min in a Beckman TLA100 rotor. After centrifugation, supernatants were removed, and pellets were resuspended in an equal volume of buffer. Samples were then subjected to SDS-PAGE and visualized by Coomassie Blue staining. For tubulation assays, 3 μM wt AMPH-1 or 15 μM ΔH_0_ were incubated with 60 pM, 200 nm PS liposomes for 20 min, prior to grid application and staining. Samples were applied to formvar/carbon-coated 400 mesh copper grids (Electron Microscopy Services) and stained with 2% uranyl acetate. The grids were imaged using a JEOL 1200EX transmission electron microscope at 100 kV.

### Disulfide Cross-linking

Oxidative disulfide crosslinking was carried out in the presence of 1 mM copper (II)-(1,10-phenanthroline)_3_ (CuPh). In all cases, a 10 mM CuPh stock solution was prepared for each experiment by mixing 20 mM 1,10-phenanthroline with 300 mM CuSO_4_ in 20% ethanol immediately prior to use. For experiments in the absence of membrane, AMPH-1 cysteine mutants (5 µM) were first subjected to ultracentrifugation, as noted above, to remove a small population of oligomers that form during purification and storage. Protein samples were then incubated in either the presence of absence of 500 µM GTP, GDP, or GMPPNP for 60 min at 23 °C, followed by addition of 1 mM CuPh and incubation for 5 min at 23 °C. The reaction was quenched with 10 mM EDTA and samples were analyzed using non-reducing SDS-PAGE and Coomassie Blue. In the presence of membranes, samples of AMPH-1 cysteine mutants (10 µM) were first clarified using ultracentrifugation as noted above and then incubated with 0.3 mg/ml lipids in the presence or absence of 500 µM GTP, GDP, or GMPPNP for 20 min. Following addition of 1 mM CuPh, samples were incubated for 2 min at 23 °C then treated with 10 mM EDTA. Samples were centrifuged at 90,000x g for 15 min in a TLA100 rotor using an Optima MAX-XP ultracentrifuge (Beckman). Pellets were re-suspended and analyzed using non-reducing SDS-PAGE and Coomassie Blue staining.

### Förster Resonance Energy Transfer (FRET)

FRET experiments were conducted using a T-format steady-state fluorometer (Photon Technologies International) at a temperature of 23 °C. The donor fluorophore, EDANS, was excited at a wavelength of 336 ± 2 nm, and emission signals for both the donor and acceptor (fluorescein) were recorded between 400 - 600 nm using a bandwidth of ± 2 nm. For experiments conducted in the absence of membrane, 100 nM dual-labeled AMPH-1 was first mixed with 10 µM unlabeled wt AMPH-1 to inhibit inter-dimer FRET coupling. This mixture was incubated with 1 mM GDP or GMPPNP (Sigma-Aldrich) for 1 min prior to fluorescence spectra acquisition. Intra-monomer FRET coupling was established using samples of double-labeled AMPH-1 treated with 100 ng/ml trypsin for 5 hours prior (Figure S3). For experiments conducted in the presence of liposomes, dual-labeled AMPH-1 (100 nM) was first mixed with 0.3 mg/mL 200 nm PS liposomes, supplemented with either 1 mM GDP or GMPPNP (Sigma-Aldrich) and incubated for 2 minutes at 23 °C. Unlabeled wt AMPH-1 was then added to a concentration of 10 µM and the sample was incubated for an additional 20 minutes prior to fluorescence data collection. The increased scattering of liposome-containing samples was corrected for by subtracting the spectral response of 10.1 µM unlabeled wt AMPH-1 in the presence of 0.3 mg/mL 200 nm PS liposomes from the emission spectra of samples containing fluorescent AMPH-1 proteins.

### Membrane Penetration Assays

AMPH-1 C49A/C110S/F5C was labeled at amino acid position 5 using IANBD, as described above. The resulting NBD-AMPH-1 was mixed with 0.3 mg/mL 200 nm liposomes created from either 100% PS, 90% PS 10% PC, or 90% PS 10% 5-doxyl-PC, in the presence or absence of 500 µM GDP or GMPPNP. In all cases, liposome samples were first incubated with NBD-AMPH-1, with or without nucleotides, for 2 minutes at 23 °C prior to the addition of 10 µM wt AMPH-1. Sample were incubated for an additional 20 min at 23 °C prior to collection of fluorescence emission spectra between 500-600 nm (2 nm bandpass) with excitation at 480 nm ± 2 nm. The spectral response of matched samples containing only wt AMPH-1 at final concentration of 10.1 µM was used to correct for the enhanced scatter introduced by the liposomes.

### Cryo-EM sample preparation and data acquisition

AMPH-1-bound membrane tubules were prepared by first incubating wt AMPH-1 (30 µM) with 1 µm diameter liposomes (0.6 mg/mL) and 1 mM GTP for 60 min at room temperature. The mixture was applied to glow-discharged holey carbon grid (Quantifoil, R2/1, 300 mesh) mounted in a Vitrobot Mark IV (ThermoFisher), incubated for 5 min at 10 °C at 100% relative humidity, followed by blotting (4 s) and vitrification in liquid ethane. Cryo-EM images were recorded using a Titan Krios G4 microscope (ThermoFisher) at 300 kV.

Automated data acquisition was achieved on a Gatan K3 direct detection camera (Gatan) with a pixel size of 1.07Å and defocus ranged from -0.5 to -2.5 μm. The beam intensity was adjusted to a total electron dose of 60 e−/Å^2^ for a 40-frames movie stack. A total of 9,662 movie stacks were recorded.

### Cryo-EM data processing and structure determination

CryoEM micrographs were initially processed using cryoSPARC v.4.0 ^72^, and the overall processing pipeline employed is illustrated in Figure S5. Motion correction and contrast transfer function determinations were performed using the Motion Correction and CTF patches. Application of iterative filament tracer following curate exposures produced 4,066 micrographs containing well-defined tubules. Two-dimensional (2D) classification was then used to generate a dataset of 629,541 tubule sub-particles of pixel size 4.27 Å. These particles were subjected to iterative 2D classification until visible structural features were observed. The clearest sub-class yielded 43,581 particles and was selected for additional processing. A pair of self-calculated helical rise and twist parameters was used during symmetry searching to efficiently narrow the search range. The predicted rise and twist parameters were then subjected to helical refinement until a reasonable map was generated.

### Atomic model building and refinement

To build the model of all structures in this study, initial models were predicted by Alphafold2 ^51^ and manually docked into our cryo-EM map. Iterative manual adjustment and rebuilding of models were done with ISOLDE ^73^, which was followed by the real-space refinement in PHENIX ^74^. All structural figures were prepared using UCSF ChimeraX ^75^.

## Acknowledgements

This research has been supported by the National Institutes of Health grant (H.R. GM114405 and GM134063). Y.W. and J.Z. were supported by a grants from the National Institutes of Health (J.Z. R01GM141659), National Science Foundation (J.Z. MCB-1902392), and a TPT grant from Texas A&M University. We would like to thank Dr. Gaya Yadav, technical director of the Laboratory for Biomolecular Structure and Dynamics at Texas A&M University for his assistance in the collection and curation of cryoEM data sets. We would also like to thank Dr. Jirapat Thongchol for discussions about cryoEM data processing and members of the Rye lab for their valuable suggestions on this manuscript.

## Author Statement

**Wei Gai**: Investigation, Formal Analysis, Writing - Review and Editing; **Yuang Wang**: Investigation, Validation, Formal Analysis; **Junjie Zhang**: Formal Analysis, Supervision, Writing - Review and Editing,; **Chavela Carr**: Conceptualization, Writing - Original Draft, Writing - Review and Editing, Supervision; **Hays S. Rye**: Conceptualization, Methodology, Writing - Original Draft, Writing - Review and Editing, Supervision, Funding Acquisition.

## Ethics Declaration

The authors declare that they have no conflict of interest.

## Data Availability Statement

The data that support the findings of this study are available upon reasonable request from the corresponding author. The coordinate and cryo-EM map are deposited in the Protein Data Bank (PDB) and the Electron Microscopy Data Bank (EMDB) with the accession codes: 9BQX and EMD-44828.

---

**Figure S1.**
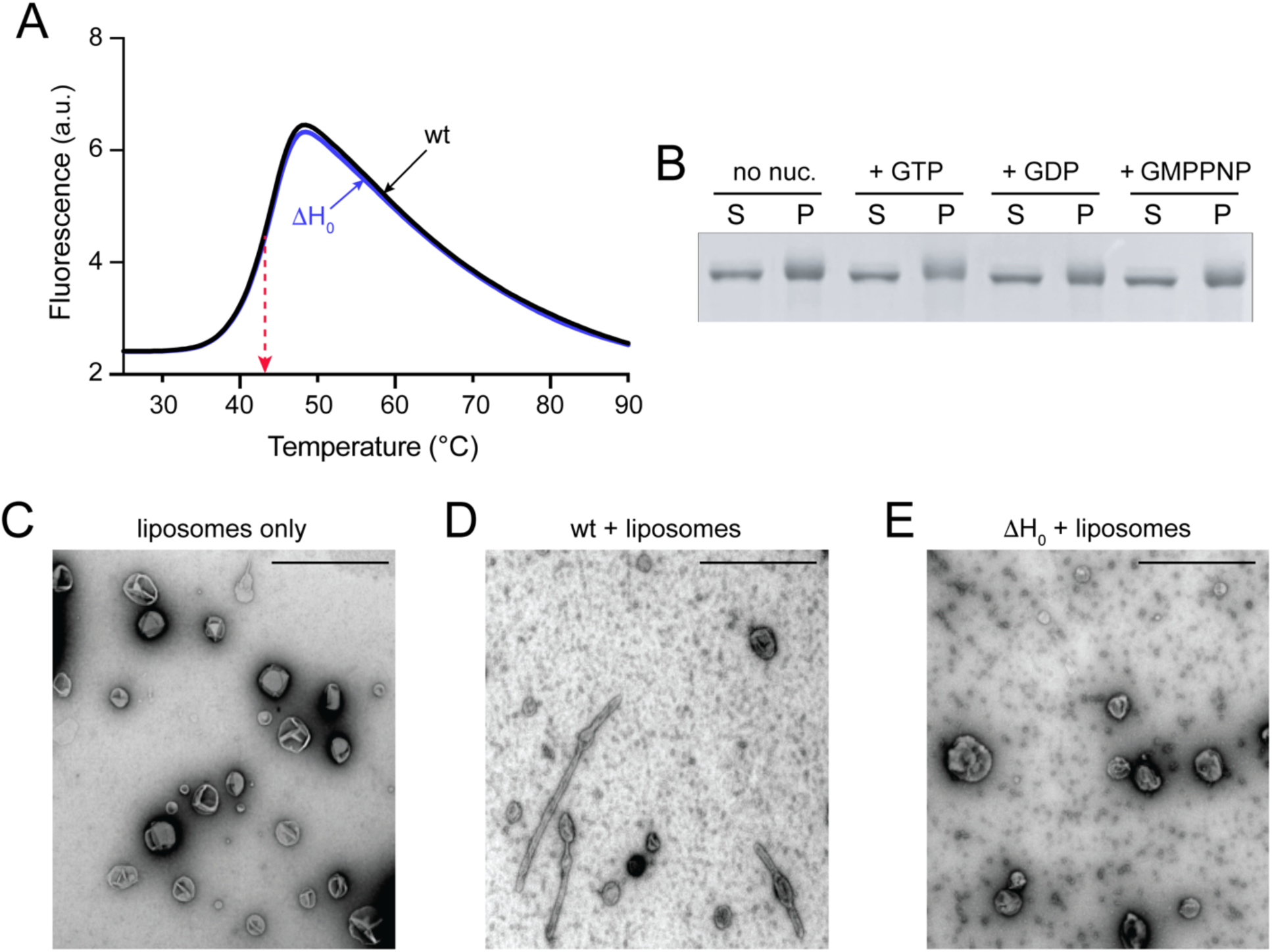
Removal of the H_0_ helix blocks membrane tubulation but has no impact on AMPH-1 dimer stability. (A) Fluorescence thermal shift assay of wild-type AMPH-1 (wt) and the AMPH-1 H_0_ truncation variant (ΔH_0_). Samples of each protein were mixed (1 µM, final concentration) with SYPRO Orange (the final concentration obtained by diluting a 5000x stock solution to 5x) in 25 mM Tris, pH 7.4, 150 mM KCl, 1 mM MgCl_2_, 1 mM DTT, and subjected to a thermal ramp from 23 °C to 95 °C in 0.5 °C increments, using a 10 s equilibration dwell at each temperature. The apparent T_m_ of the proteins is the same (red arrow). (B) Coomassie-stained SDS-PAGE gel showing the supernatant (S) and pellet (P) fractions of an wt AMPH-1 liposome binding assay carried out in the presence of different guanine nucleotides. Liposomes (100% PS) were created by extrusion (0.33 mg/ml final, 200 nm average diameter) and mixed with either GTP, GDP, or GMPPNP (500 µM). Each liposome-nucleotide mixture was then supplemented with 2 μM wt AMPH-1 and incubated at 23 °C for 10 min prior to centrifugation (90,000x g for 15 min). The morphology of 200 nm PS liposomes after mixing with either (C) no protein (D) wt AMPH-1 or (E) AMPH-1 (ΔH_0_) was examined using negative stain electron microscopy in the absence of nucleotide. Scale bar represents 1 µm. While tubules are readily observed in the presence of wild type AMPH-1, no tubules were observed in either liposome only samples or in the presence of the ΔH_0_ truncation variant.

**Figure S2.**
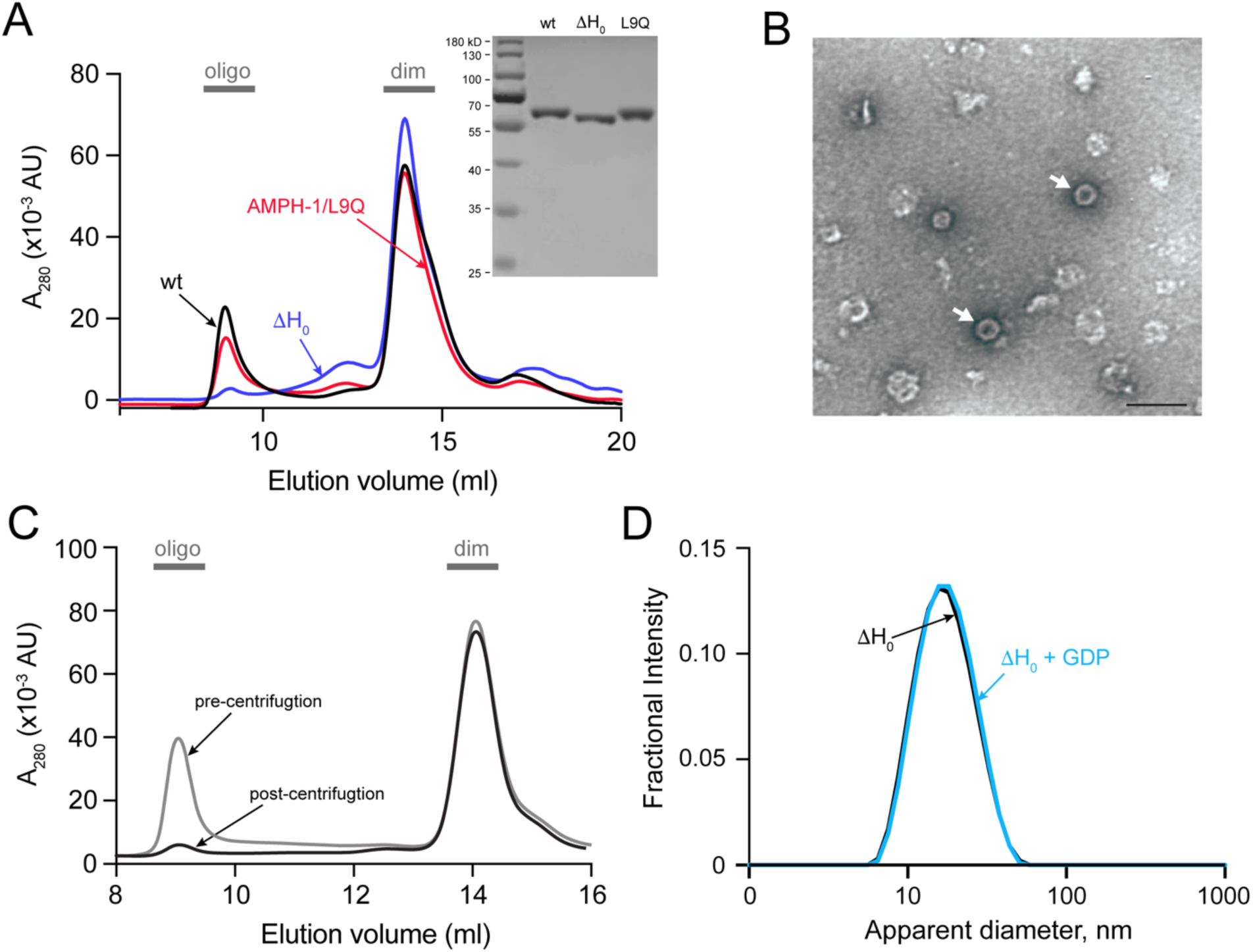
The H_0_ helix is required for formation of AMPH-1 oligomers in free solution. (A) Size-exclusion chromatography (SEC) analysis of wt, ΔH_0_ and L9Q variants of AMPH-1 results in two distinct peaks. The first peak (elution volume ≈ 9 ml) is composed of large, oligomeric assemblies of AMPH-1 (oligo), while the second peak (elution volume ≈ 14 ml) is composed of dimeric AMPH-1 (dim). The inset shows a Coomassie-stained SDS-PAGE gel of each protein sample used for the SEC analysis. (B) Examination of the wt AMPH-1 oligomer peak by negative-stain EM reveals a range of assembled AMPH-1 complexes, including a distinct, ring-like oligomer (indicated by arrows) possessing an apparent diameter of 50 ± 2 nm. (C) The SEC profile of wt AMPH-1 before and after ultracentrifugation (300,000x g for 30 min) shows a loss of the early eluting peak by selective sedimentation of higher molecular weight oligomeric species. (D) The dynamic light scattering (DLS) profiles of the ΔH_0_ AMPH-1 variant, in the presence and absence of GDP (500 µM), shows no detectable formation of higher molecular weight particles following incubation at 23 °C for 1 hr.

**Figure S3.**
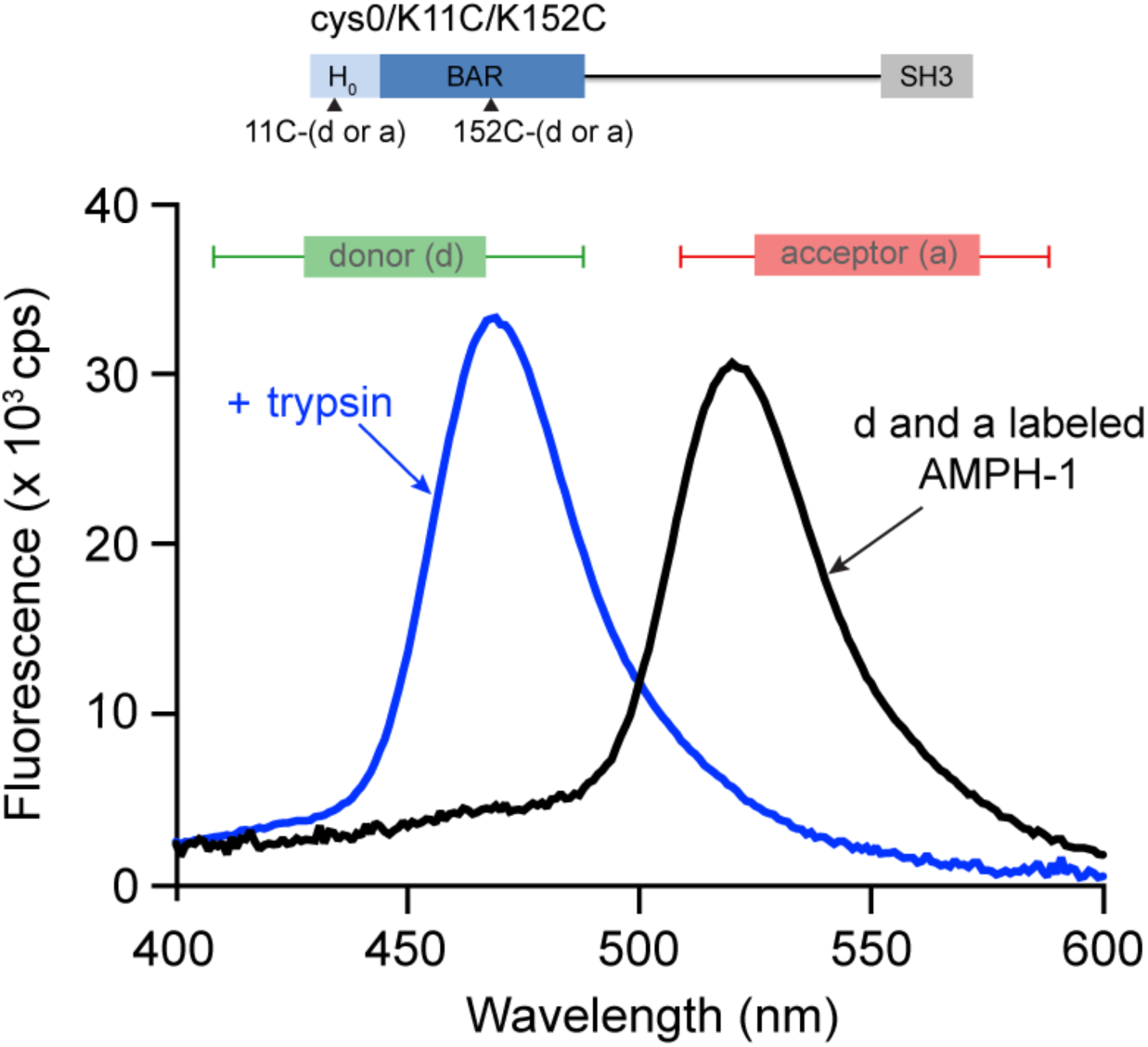
Protease digestion demonstrates highly efficient FRET coupling between labeled K11C and K152C sites. Fluorescence emission spectra for double labeled (donor and acceptor) AMPH-1 are shown before (black) and after (blue) trypsinolysis. A sample of double-labeled AMPH-1 mixed with a large excess of unlabeled AMPH-1 was prepared by mixing 0.5 µM double-labeled AMPH-1 with 10 µM unlabeled wt AMPH-1. An aliquot of this sample was treated with 100 ng/ml of trypsin for 5 hr at 23 °C. The emission spectra for both the untreated and trypsinized samples were collected with excitation at the EDANS donor absorption maximum (336 nm ± 5 nm). The near total quenching of the EDANS donor and large emission from the fluorescein acceptor in the intact protein, compared to the dramatic de-quenching of the donor and near total loss of sensitized fluorescence from the acceptor following trypsinolysis, demonstrates the highly efficient FRET coupling between the probes in the native AMPH-1 dimer.

**Figure S4.**
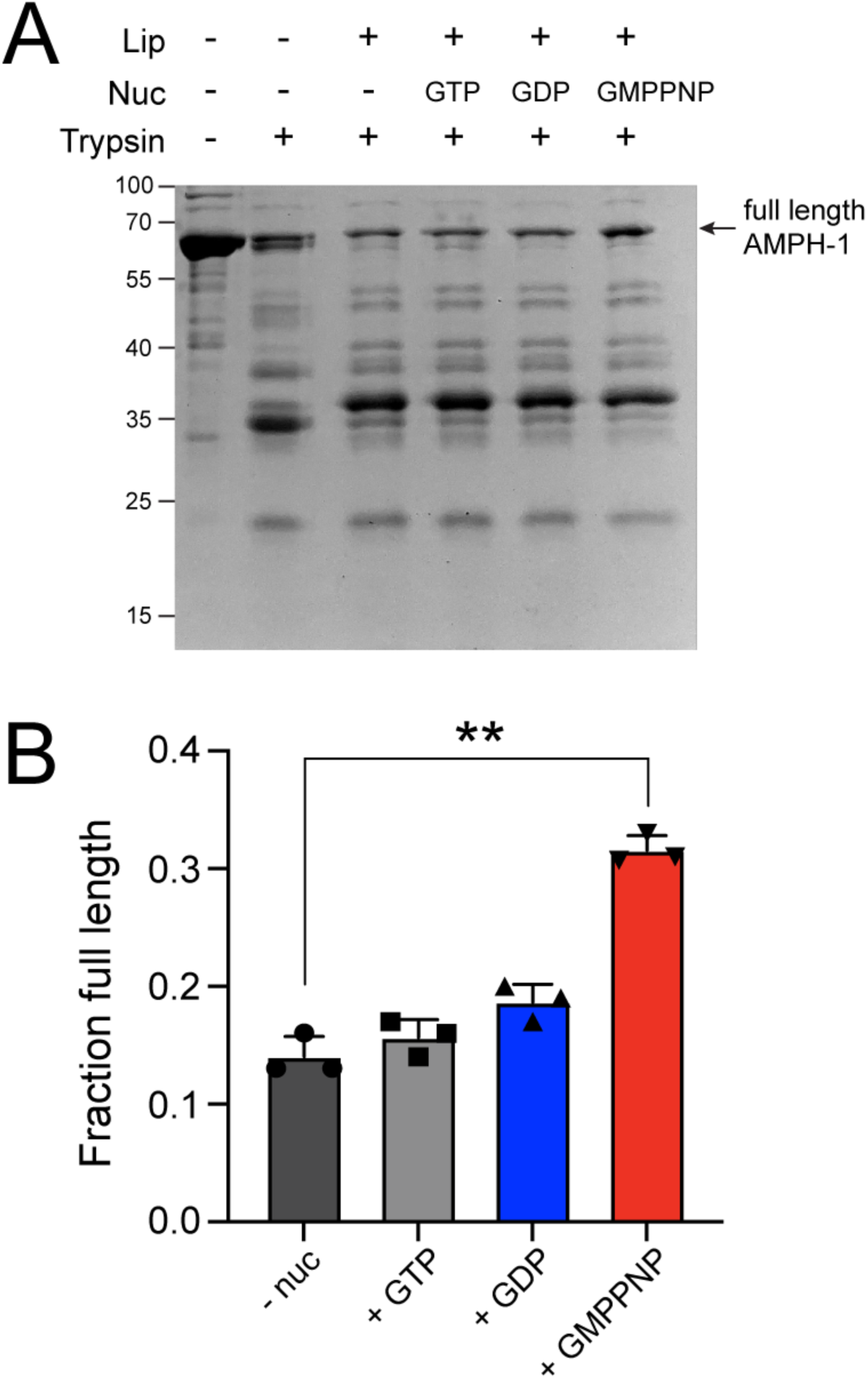
Non-hydrolysable GMPPNP increases the protease resistance of membrane-bound AMPH-1. (A) Membrane-bound AMPH-1 was treated with trypsin in the presence and absence of guanine nucleotides. Following centrifugation (300,000x g for 30 min), wt AMPH-1 (2 μM) was incubated for 20 min at 23 °C either alone, with 200 nm, 100% PS liposomes (0.15 mg/ml) or with liposomes and 500 µM GTP, GDP, or GMPPNP. Samples were then treated with trypsin (1 ng/µl) for 10 minutes, quenched with PMSF (1 mM) and analyzed by SDS-PAGE. (B) The amount of full length AMPH-1 remaining was quantified by densitometry and normalized to samples not treated with trypsin. The plot shows the mean and error bars (± s.d) of n = 3 independent experimental replicates. Significance was assessed with a one-way ANOVA using the Geyser-Greenhouse correction (**P = 0.004).

**Figure S5.**
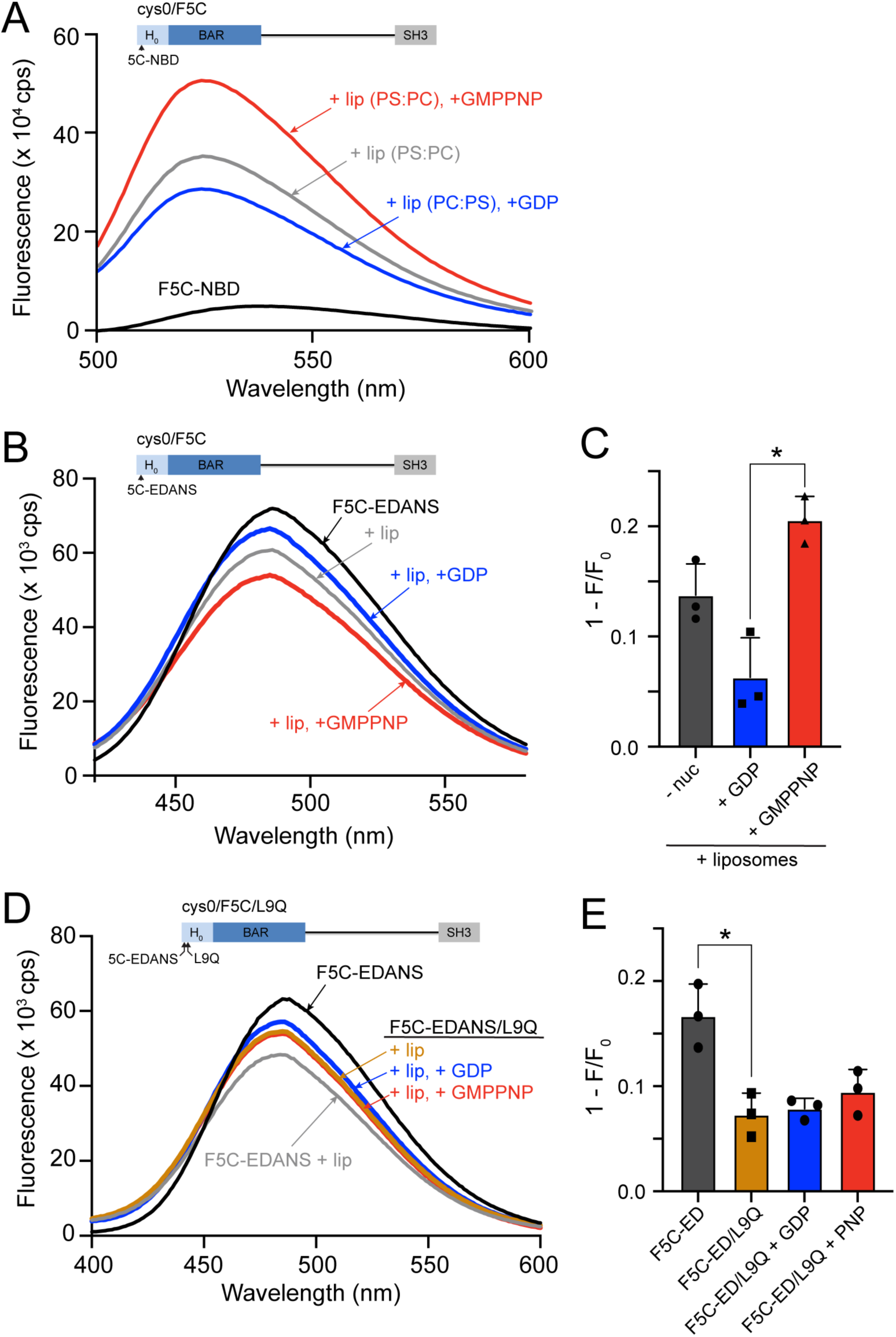
GDP stabilizes a shallower penetration of the H0 helix into the lipid bilayer. (A) Liposomes containing 0.3 mg/mL with a diameter of 200 nm, consisting of 90% PS and 10% PC, were initially treated with 100 nM NBD-AMPH-1 for 2 minutes. Subsequently, unlabeled wt AMPH-1 was introduced at a concentration of 10 µM, and the mixture was further incubated for 20 minutes in the presence or absence of GDP (500 µM) or GMPPNP (500 µM). Fluorescence emission spectra from 500 nm to 600 nm were recorded using an excitation wavelength of 480 nm, with background fluorescence subtracted as described. (B) Domain structure of AMPH-1 cys0 with a single Cys incorporated into the H_0_ helix (F5C) that was labeled with the fluorescent dye IAEDANS (wt-cys0/F5C-EDANS). Steady state fluorescence emission spectra (lambda ex = 336 ± 2 nm) for wt-cys0/F5C-EDANS (labeled protein 0.1 µM, unlabeled wt AMPH-1 10 µM) in the absence (black) and presence of 200 nm, 100% PS liposomes (0.3 mg/mL; gray). The quenching of the EDANS probe upon addition of liposomes is indicative of a change in the local environment of the EDANS probe caused by partial membrane insertion of this face of the H_0_ helix ^50^. Addition of GDP (500 µM; blue) results in de-quenching of the EDANS probe, consistent with a reduction in average helical penetration into the membrane. The presence of GMPPNP (500 µM, red) results in an increase in quenching, consistent with an increase in average helical penetration. (C) Quantitation of EDANS fluorescence quenching (1-F/F_0_) with plot showing the mean and error bars (± s.d) of n = 3 independent experimental replicates. Significance was assessed with a one-way ANOVA using the Geyser-Greenhouse correction with *P = 0.0282. (D) Domain structure of wt-cys0/F5C-EDANS to which the inhibitory mutation (L9Q) has been added wt-cys0/F5C-EDANS/L9Q. Steady state fluorescence emission spectra (lambda ex = 336 ± 2 nm) for wt-cys0/F5C-EDANS/L9Q (labeled protein 0.1 µM, unlabeled wt AMPH-1 10µM) in the presence of PS liposomes alone (0.3 mg/mL; orange) or liposomes and 500 µM GDP (blue) or GMPPNP (red). The emission of wt-cys0/F5C-EDANS/L9Q in the absence of liposomes (not shown) is identical to that of wt-cys0/F5C-EDANS (black). The quenching of the EDANS probe upon addition of liposomes for wt-cys0/F5C-EDANS (gray) is shown for reference. Addition of the inhibitory L9Q mutation both reduces the level of EDANS quenching upon liposome addition and displays no significant dependence on the presence of guanine nucleotides. (E) Quantitation of membrane-induced EDANS quenching for wt-cys0/F5C-EDANS compared to wt-cys0/F5C-EDANS/L9Q with the plot showing the mean and error bars (± s.d) of n = 3 independent experimental replicates. Significance was assessed with a one-way ANOVA using the Geyser-Greenhouse correction with *P = 0.0447.

**Figure S6.**
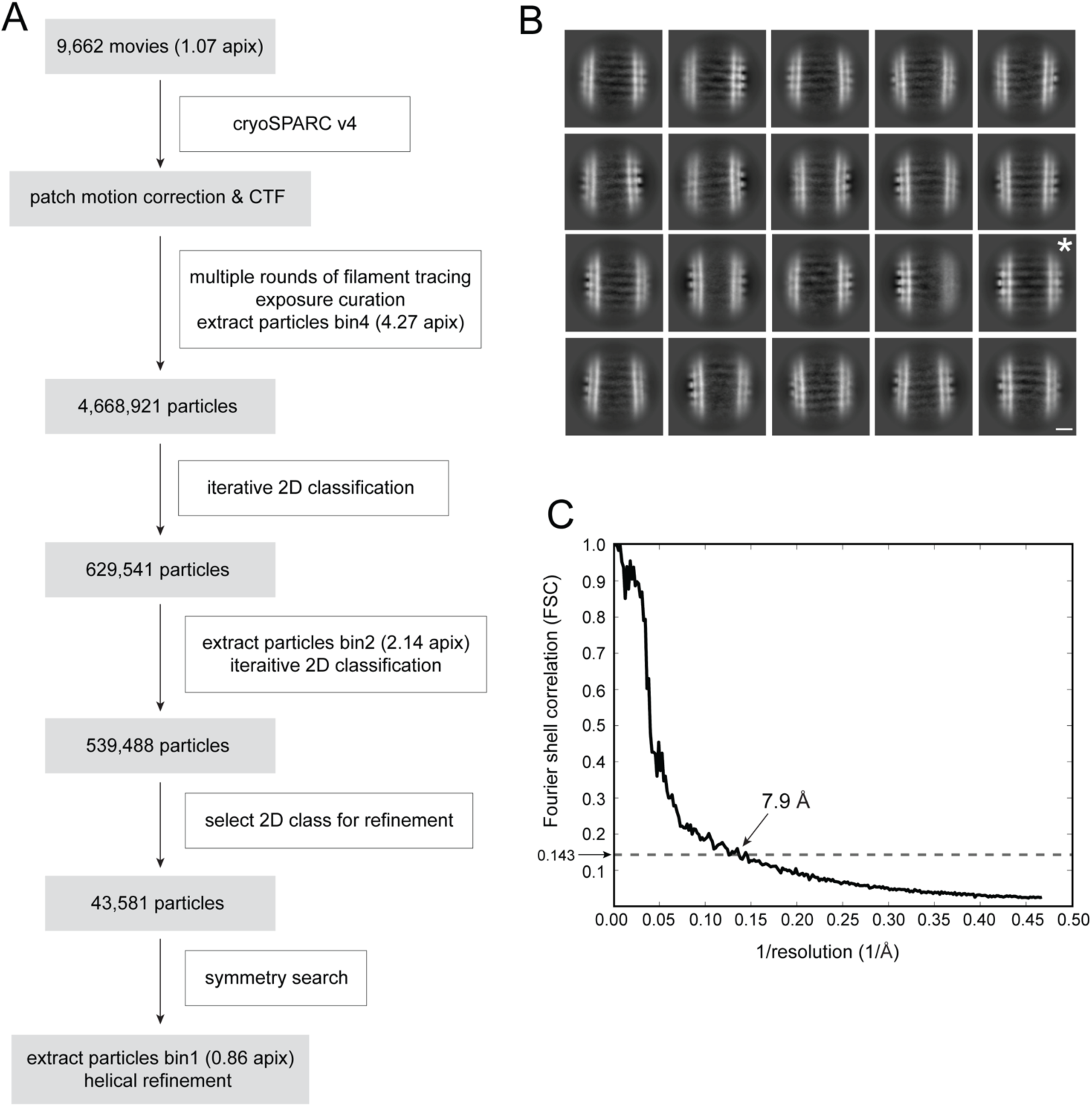
CryoEM data processing and refinement pipeline. (A) Cryo-EM data processing diagram of the AMPH-1 bound membrane tubules incubated with 1 mM GTP for one hour. (B) Representative 2D class averages of the tube. The selected class for refinement is indicated by white asterisk (*). The white scale bar denotes 100 Å. (C) The gold-standard Fourier Shell Correlation (FSC) curves of the density map of the tubule.

## References

1. Grant, B. D. & Donaldson, J. G. Pathways and mechanisms of endocytic recycling. Nat Rev Mol Cell Bio 10, 597–608 (2009).

2. Naslavsky, N. & Caplan, S. The enigmatic endosome - sorting the ins and outs of endocytic trafficking. J Cell Sci 131, jcs216499 (2018).

3. Campelo, F. & Malhotra, V. Membrane Fission: The Biogenesis of Transport Carriers. Annual review of biochemistry 81, 407–427 (2012).

4. Chen, Y., Huang, L., Qi, X. & Chen, C. Insulin Receptor Trafficking: Consequences for Insulin Sensitivity and Diabetes. Int. J. Mol. Sci. 20, 5007 (2019).

5. Coster, A. C. F., Govers, R. & James, D. E. Insulin Stimulates the Entry of GLUT4 into the Endosomal Recycling Pathway by a Quantal Mechanism. Traffic 5, 763–771 (2004).

6. O’Sullivan, M. J. & Lindsay, A. J. The Endosomal Recycling Pathway—At the Crossroads of the Cell. Int J Mol Sci 21, 6074 (2020).

7. Burr, M. L. et al. CMTM6 maintains the expression of PD-L1 and regulates anti-tumour immunity. Nature 549, 101–105 (2017).

8. Mishra, A., Hourigan, D. & Lindsay, A. J. Inhibition of the endosomal recycling pathway downregulates HER2 activation and overcomes resistance to tyrosine kinase inhibitors in HER2-positive breast cancer. Cancer Lett. 529, 153–167 (2022).

9. Gleason, R. J., Akintobi, A. M., Grant, B. D. & Padgett, R. W. BMP signaling requires retromer-dependent recycling of the type I receptor. Proc. Natl. Acad. Sci. 111, 2578– 2583 (2014).

10. Li, X. & DiFiglia, M. The recycling endosome and its role in neurological disorders. Prog. Neurobiol. 97, 127–141 (2012).

11. Antonny, B. et al. Membrane fission by dynamin: what we know and what we need to know. The EMBO journal 35, 2270–2284 (2016).

12. Renard, H.-F., Johannes, L. & Morsomme, P. Increasing Diversity of Biological Membrane Fission Mechanisms. Trends in Cell Biology 1–13 (2018) doi:10.1016/j.tcb.2017.12.001.

13. Itoh, T. & Camilli, P. D. BAR, F-BAR (EFC) and ENTH/ANTH domains in the regulation of membrane-cytosol interfaces and membrane curvature. Biochim. Biophys. acta 1761, 897–912 (2006).

14. Daumke, O. et al. Architectural and mechanistic insights into an EHD ATPase involved in membrane remodelling. Nature 449, 923–927 (2007).

15. Deo, R. et al. ATP-dependent membrane remodeling links EHD1 functions to endocytic recycling. Nat Commun 9, 5187 (2018).

16. Hoernke, M. et al. EHD2 restrains dynamics of caveolae by an ATP-dependent, membrane-bound, open conformation. Proc Natl Acad Sci U S A 201614066–10 (2017) doi:10.1073/pnas.1614066114.

17. Kustigian, L. et al. GTP-stimulated membrane fission by the N-BAR protein AMPH-1. Traffic 24, 34–47 (2023).

18. Masuda, M. et al. Endophilin BAR domain drives membrane curvature by two newly identified structure-based mechanisms. EMBO J. 25, 2889–2897 (2006).

19. Brooks, A. et al. Single particle fluorescence burst analysis of epsin induced membrane fission. PloS one 10, e0119563 (2015).

20. Zhukovsky, M. A., Filograna, A., Luini, A., Corda, D. & Valente, C. Protein Amphipathic Helix Insertion: A Mechanism to Induce Membrane Fission. Frontiers Cell Dev Biology 7, 291 (2019).

21. Peter, B. J. et al. BAR domains as sensors of membrane curvature: the amphiphysin BAR structure. Science 303, 495–499 (2004).

22. Gallop, J. L. et al. Mechanism of endophilin N-BAR domain-mediated membrane curvature. The EMBO J. 25, 2898–2910 (2006).

23. Bhatia, V. K. et al. Amphipathic motifs in BAR domains are essential for membrane curvature sensing. EMBO J. 28, 3303–3314 (2009).

24. Drin, G. & Antonny, B. Amphipathic helices and membrane curvature. FEBS Lett. 584, 1840–1847 (2010).

25. Pant, S. et al. AMPH-1/Amphiphysin/Bin1 functions with RME-1/Ehd1 in endocytic recycling. Nature cell biology 11, 1399–1410 (2009).

26. McMahon, H. T. & Gallop, J. L. Membrane curvature and mechanisms of dynamic cell membrane remodelling. Nature 438, 590–6 (2005).

27. Grant, B. et al. Evidence that RME-1, a conserved C. elegans EH-domain protein, functions in endocytic recycling. Nature cell biology 3, 573–579 (2001).

28. Shi, A. et al. A Novel Requirement for C. elegans Alix/ALX-1 in RME-1-Mediated Membrane Transport. Curr. Biol. 17, 1913–1924 (2007).

29. McMahon, H. T. & Boucrot, E. Membrane curvature at a glance. Journal of Cell Science 128, 1065–1070 (2015).

30. Bassereau, P. et al. The 2018 biomembrane curvature and remodeling roadmap. J Phys D Appl Phys 51, 343001 (2018).

31. Boucrot, E. et al. Membrane Fission Is Promoted by Insertion of Amphipathic Helices and Is Restricted by Crescent BAR Domains. Cell 149, 124–136 (2012).

32. Campelo, F., McMahon, H. T. & Kozlov, M. M. The Hydrophobic Insertion Mechanism of Membrane Curvature Generation by Proteins. Biophys. J. 95, 2325–2339 (2008).

33. Schmidt, N. W. & Wong, G. C. L. Antimicrobial peptides and induced membrane curvature: Geometry, coordination chemistry, and molecular engineering. Curr. Opin. Solid State Mater. Sci. 17, 151–163 (2013).

34. Zemel, A., Ben-Shaul, A. & May, S. Modulation of the Spontaneous Curvature and Bending Rigidity of Lipid Membranes by Interfacially Adsorbed Amphipathic Peptides. J. Phys. Chem. B 112, 6988–6996 (2008).

35. Frost, A., Unger, V. M. & Camilli, P. D. The BAR Domain Superfamily: Membrane-Molding Macromolecules. Cell 137, 191–196 (2009).

36. Mim, C. et al. Structural Basis of Membrane Bending by the N-BAR Protein Endophilin. Cell 149, 137–145 (2012).

37. Bhatt, V. S., Ashley, R. & Sundborger-Lunna, A. Amphipathic Motifs Regulate N-BAR Protein Endophilin B1 Auto-inhibition and Drive Membrane Remodeling. Structure (2020) doi:10.1016/j.str.2020.09.012.

38. Bonazzi, F. & Weikl, T. R. Membrane Morphologies Induced by Arc-Shaped Scaffolds Are Determined by Arc Angle and Coverage. Biophys. J. 116, 1239–1247 (2019).

39. Simunovic, M. et al. How curvature-generating proteins build scaffolds on membrane nanotubes. Proc. Natl. Acad. Sci. 113, 11226–11231 (2016).

40. Noguchi, H. Membrane tubule formation by banana-shaped proteins with or without transient network structure. Sci. Rep. 6, 20935 (2016).

41. Liu, J., Kaksonen, M., Drubin, D. G. & Oster, G. Endocytic vesicle scission by lipid phase boundary forces. Proc Natl Acad Sci U S A 103, 10277–10282 (2006).

42. Renard, H.-F. et al. Endophilin-A2 functions in membrane scission in clathrin-independent endocytosis. Nature 517, 493–496 (2015).

43. Day, C. A. et al. Microtubule Motors Power Plasma Membrane Tubulation in Clathrin-Independent Endocytosis. Traffic 16, 572–590 (2015).

44. Simunovic, M. et al. Friction Mediates Scission of Tubular Membranes Scaffolded by BAR Proteins. Cell 170, 172–184.e11 (2017).

45. Chen, Z., Zhu, C., Kuo, C. J., Robustelli, J. & Baumgart, T. The N-Terminal Amphipathic Helix of Endophilin Does Not Contribute to Its Molecular Curvature Generation Capacity.J. Am. Chem. Soc. 138, 14616–14622 (2016).

46. Kyrychenko, A., Rodnin, M. V. & Ladokhin, A. S. Calibration of Distribution Analysis of the Depth of Membrane Penetration Using Simulations and Depth-Dependent Fluorescence Quenching. J. Membr. Biol. 248, 583–594 (2015).

47. Chattopadhyay, A. & London, E. Parallax method for direct measurement of membrane penetration depth utilizing fluorescence quenching by spin-labeled phospholipids. Biochemistry 26, 39–45 (1987).

48. London, E. & Ladokhin, A. S. Measuring the depth of amino acid residues in membrane-inserted peptides by fluorescence quenching. Curr. Top. Membr. 52, 89–115 (2002).

49. Jiang, C. et al. NBD-based synthetic probes for sensing small molecules and proteins: design, sensing mechanisms and biological applications. Chem. Soc. Rev. 50, 7436– 7495 (2021).

50. Nair, M. S. & Dean, D. H. All domains of Cry1A toxins insert into insect brush border membranes. J. Biol. Chem. 283, 26324–31 (2008).

51. Jumper, J. et al. Highly accurate protein structure prediction with AlphaFold. Nature 596, 583–589 (2021).

52. Isas, J. M., Ambroso, M. R., Hegde, P. B., Langen, J. & Langen, R. Tubulation by Amphiphysin Requires Concentration-Dependent Switching from Wedging to Scaffolding. Structure/Folding and Design 23, 1–10 (2015).

53. Cui, H. et al. Understanding the Role of Amphipathic Helices in N-BAR Domain Driven Membrane Remodeling. Biophys. J. 104, 404–411 (2013).

54. Jao, C. C. et al. Roles of Amphipathic Helices and the Bin/Amphiphysin/Rvs (BAR) Domain of Endophilin in Membrane Curvature Generation*. J. Biol. Chem. 285, 20164– 20170 (2010).

55. Ambroso, M. R., Hegde, B. G. & Langen, R. Endophilin A1 induces different membrane shapes using a conformational switch that is regulated by phosphorylation. Proceedings of the National Academy of Sciences of the United States of America 111, 6982–6987 (2014).

56. Schmidt, N. W. & Wong, G. C. L. Antimicrobial peptides and induced membrane curvature: Geometry, coordination chemistry, and molecular engineering. Curr. Opin. Solid State Mater. Sci. 17, 151–163 (2013).

57. Carman, P. J. & Dominguez, R. BAR domain proteins—a linkage between cellular membranes, signaling pathways, and the actin cytoskeleton. Biophys. Rev. 10, 1587– 1604 (2018).

58. Qualmann, B., Koch, D. & Kessels, M. M. Let’s go bananas: revisiting the endocytic BAR code. Embo J 30, 3501–3515 (2011).

59. Hanna, M. G. et al. Sar1 GTPase Activity Is Regulated by Membrane Curvature* ♦. J. Biol. Chem. 291, 1014–1027 (2016).

60. Lundmark, R., Doherty, G. J., Vallis, Y., Peter, B. J. & McMahon, H. T. Arf family GTP loading is activated by, and generates, positive membrane curvature. Biochem. J. 414, 189–194 (2008).

61. Bonazzi, F., Hall, C. K. & Weikl, T. R. Membrane morphologies induced by mixtures of arc-shaped particles with opposite curvature. Soft Matter 17, 268–275 (2020).

62. Gao, J., Hou, R., Hu, W., Weikl, T. R. & Hu, J. Which Coverages of Arc-Shaped Proteins Are Required for Membrane Tubulation? J. Phys. Chem. B 128, 4735–4740 (2024).

63. Yoon, Y., Zhang, X. & Cho, W. Phosphatidylinositol 4,5-Bisphosphate (PtdIns(4,5)P2) Specifically Induces Membrane Penetration and Deformation by Bin/Amphiphysin/Rvs (BAR) Domains. 287, 34078–34090 (2012).

64. Wu, T. & Baumgart, T. BIN1 Membrane Curvature Sensing and Generation Show Autoinhibition Regulated by Downstream Ligands and PI(4,5)P2. Biochemistry-us 53, 7297–7309 (2014).

65. Adam, J., Basnet, N. & Mizuno, N. Structural insights into the cooperative remodeling of membranes by amphiphysin/BIN1. Sci. Rep. 5, 15452 (2015).

66. Aspenström, P. The Intrinsic GDP/GTP Exchange Activities of Cdc42 and Rac1 Are Critical Determinants for Their Specific Effects on Mobilization of the Actin Filament System. Cells 8, 759 (2019).

67. Kessels, M. M. & Qualmann, B. Different functional modes of BAR domain proteins in formation and plasticity of mammalian postsynapses. J. Cell Sci. 128, 3177–85 (2015).

68. Curchoe, C. L. & Manor, U. Actin Cytoskeleton-Mediated Constriction of Membrane Organelles via Endoplasmic Reticulum Scaffolding. ACS Biomater. Sci. Eng. 3, 2727–2732 (2017).

69. Rodriguez-Polanco, W. R., Norris, A., Velasco, A. B., Gleason, A. M. & Grant, B. D. Syndapin and GTPase RAP-1 control endocytic recycling via RHO-1 and non-muscle myosin II. Curr. Biol. 33, 4844–4856.e5 (2023).

70. Machesky, L. M. Rab11FIP proteins link endocytic recycling vesicles for cytoskeletal transport and tethering. Biosci. Rep. 39, BSR20182219 (2019).

71. Webb, M. R. A continuous spectrophotometric assay for inorganic phosphate and for measuring phosphate release kinetics in biological systems. Proc Natl Acad Sci U S A 89, 4884–4887 (1992).

72. Punjani, A., Rubinstein, J. L., Fleet, D. J. & Brubaker, M. A. cryoSPARC: algorithms for rapid unsupervised cryo-EM structure determination. Nat. Methods 14, 290–296 (2017).

73. Croll, T. I. ISOLDE: a physically realistic environment for model building into low-resolution electron-density maps. Acta Crystallogr. Sect. D: Struct. Biol. 74, 519–530 (2018).

74. Liebschner, D. et al. Macromolecular structure determination using X-rays, neutrons and electrons: recent developments in Phenix. Acta Crystallogr. Sect. D 75, 861–877 (2019).

75. Pettersen, E. et al. UCSF Chimera--a visualization system for exploratory research and analysis. J Comput Chem 25, 1605–1612 (2004).

